# Realtime morphological characterization and sorting of unlabeled viable cells using deep learning

**DOI:** 10.1101/2022.02.28.482368

**Authors:** Mahyar Salek, Nianzhen Li, Hou-Pu Chou, Kiran Saini, Andreja Jovic, Kevin B. Jacobs, Chassidy Johnson, Esther J. Lee, Christina Chang, Phuc Nguyen, Jeanette Mei, Krishna P. Pant, Amy Y. Wong-Thai, Quillan F. Smith, Stephanie Huang, Ryan Chow, Janifer Cruz, Jeff Walker, Bryan Chan, Thomas J. Musci, Euan A. Ashley, Maddison (Mahdokht) Masaeli

**Affiliations:** Deepcell Inc; 4025 Bohannon Dr., Menlo Park, CA 94025, USA; Department of Medicine, Genetics, & Biomedical Data Science, Stanford University, Stanford, CA USA

## Abstract

Phenotyping of single cells has dramatically lagged advances in molecular characterization and remains a manual, subjective, and destructive process. We introduce COSMOS, a platform for phenotyping and enrichment of cells based on deep learning interpretation of high-content morphology data in realtime. By training models on an atlas of >1.5 billion images, we demonstrate enrichment of unlabeled cells up to 33,000 fold. We apply COSMOS to multicellular tissue biopsy samples demonstrating that it can identify malignant cells with similar accuracy to molecular approaches while enriching viable cells for functional evaluation. We show high-dimensional embedding vectors of morphology generated by COSMOS without any need for complex sample pre-processing, gating, or bioinformatics capabilities, which enables discovery of cellular phenotypes, and integration of morphology into multi-dimensional analyses.

**One sentence summary:** A novel platform capable of high-throughput imaging and gently sorting cells using deep morphological assessment.

---

Technological advances in genomics and proteomics have enabled molecular profiling of single cells. Indeed, single cell characterization at the genomic, epigenomic, transcriptomic, and proteomic levels has been realized (Gawad, Koh, and Quake 2016; Schwartzman and Tanay 2015; Stegle, Teichmann, and Marioni 2015) and international collaborations are generating increasingly comprehensive cell atlases with exquisitely detailed molecular characterization of hundreds of cell types from multiple organisms (Rozenblatt-Rosen et al. 2017; Regev et al. 2017). In contrast, while cell morphology is often the gold standard for diagnosis and prognosis of many diseases and conditions, our conception of the physical form of single cells has changed little in centuries. Cell morphology characterization has not kept pace with advancements in molecular and functional characterization. This is largely due to the manual, time-sensitive, and subjective process of collecting cell morphology information and limited methods for sorting that do not perturb or damage the cells. Cytopathologists still classify cells stained with a limited number of chemical dyes using a small number of descriptive features (Alvarado-Kristensson and Rosselló 2019; Fischer 2020), and don’t have access to further separate and assess cells based on their characteristics. Despite approaches like laser capture microdissection, molecular characterization of captured single cells is limited and sorting cells with high viability remains an unrealized goal. The invention of fluorescent activated cell sorting (FACS), and mass cytometry allowed high-throughput unidimensional or multidimensional classification and sorting of cells, albeit with the prerequisite of labeling with known markers, and at most a couple of features - side scatter and forward scatter of light in flow cytometry - to assess cell physical form. Additionally, these approaches alter cells making them non-ideal for downstream characterization (Bendall et al. 2011; Bendall et al. 2012). There have been recent efforts to improve upon our capability to isolate cells based on their morphological traits (Schraivogel et al. 2022), but these approaches still rely on staining cells with fluorescent markers, which alters them. Additionally, they are limited by the number of morphological traits that can be visualized simultaneously and require heavily involved processes to define a small number of features to quantify morphology. Finally, the feature engineering approach falls short of the human expert assessment in richness and complexity.

Application of machine intelligence has led to multiple approaches to classify pathology slide images on par with human experts, including the recapitulation of immunohistochemistry signals from light microscopy alone (Rivenson et al. 2019). One group combined shallow (six-layer) convolutional neural network (CNN) classification of single cells with a sorting device to identify a small number of cell types (Nitta et al. 2018, 2020). Despite this progress, machine learning approaches for single cell analysis have been based on small data sets.

A platform that can identify, classify, and sort living cells based on morphology could greatly empower our understanding of biology at the single cell level. Specifically, a method to facilitate molecular characterization approaches downstream of sorting and enrichment of minimally perturbed cells could redefine our understanding of cell type and state while at the same time considerably reducing costs by concentrating the cells of interest to the investigator. Complex multicellular tissues, such as the tumor microenvironment, could be deconvoluted prior to the application of molecular assays rather than the conventional post hoc analyses using single cell characterization techniques.

Here, we introduce the COSMOS platform, a novel microfluidic optical device capable of high-throughput cell imaging and sorting using morphological information (**Fig. 1**). The hardware is complemented by (i) a deep inference infrastructure, (ii) a machine learning assisted human image annotation tool, (iii) an atlas of expert-annotated images of single cells called Deep Cell Atlas (DCA), and (iv) a library of pre-trained machine learning models for specific biological applications. COSMOS yields populations of cells that are label-free, viable, and minimally perturbed, allowing sorted cells to be recovered and further characterized by molecular and functional assays. Additionally, cell images can be used to generate high-dimensional morphological profiles to reveal and explore previously unrecognized heterogeneous cell populations. We demonstrate several applications including enrichment of tumor cells, and gene expression analysis of sorted cells.

**Figure 1.**
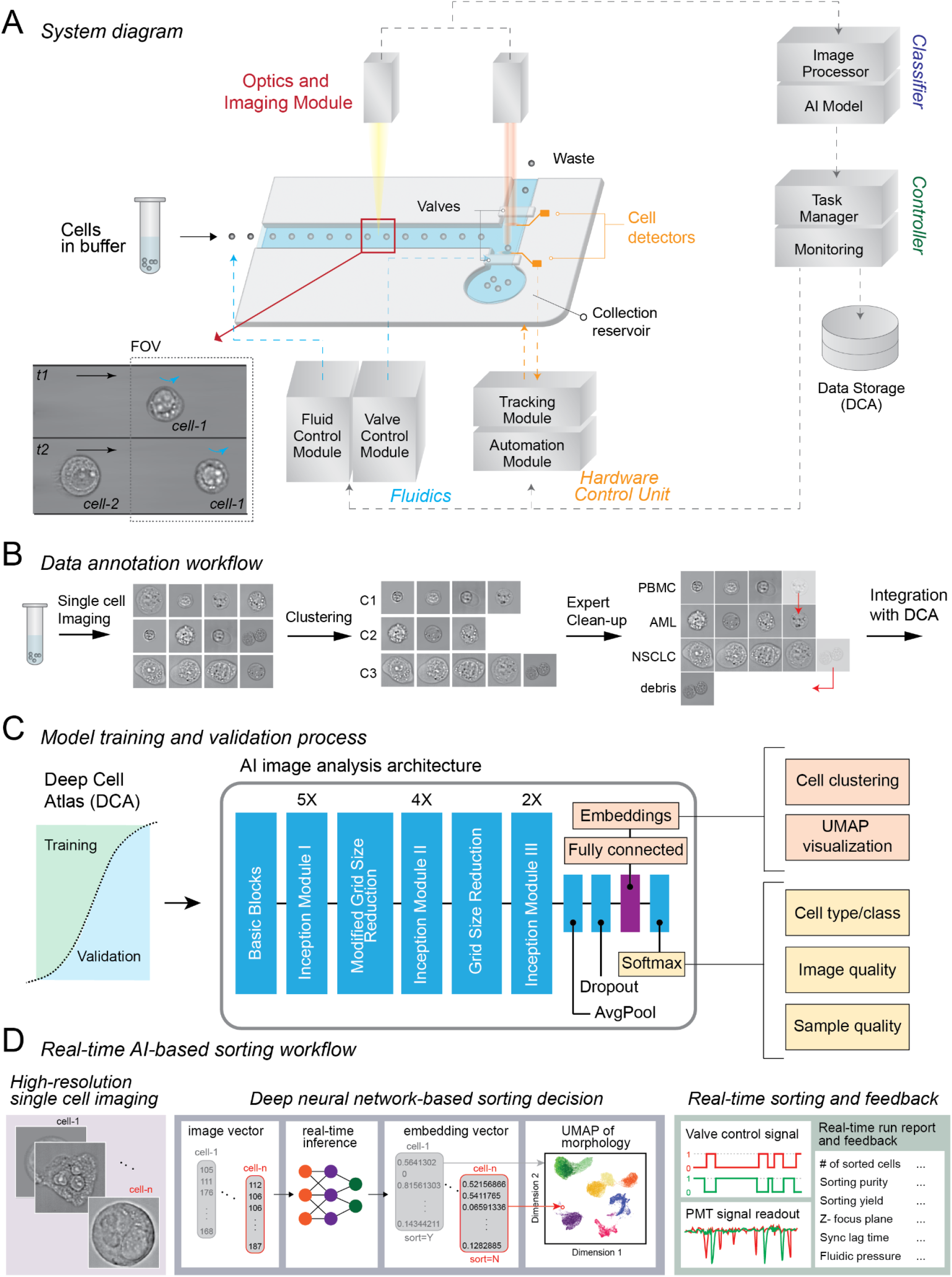
COSMOS platform schematic. **(A)** System diagram: A portion of the microfluidic cartridge and interplay between different components of the software and hardware modules are shown. Cells in suspension are inserted into the cartridge. Cells are focused on a single z plane and lateral trajectory. Two images are collected per cell. The hardware includes: i. Fluidics (Fluid Control and Valve Control Modules), ii. Optics and Imaging Module and iii. Hardware Control Unit for auto-focusing and -alignment (Tracking and Automation Modules). The software includes Classifier, Controller, and Data Storage modules. **(B)** Data annotation workflow: High contrast, bright-field images of single cells are captured while flowing in the microfluidic chip. AI-assisted image annotation software is used to cluster individual cell images. A human expert uses the labeling tool to adjust and batch-label the cell clusters. In the example shown, one acute myeloid leukemia (AML) cell was mis-clustered with a group of PBMCs and an image showing debris was mis-clustered with a group of NSCLC cells. These errors are corrected by the “Expert Clean-up” step. The annotated cells are then integrated into DCA. **(C)** Model training and validation process: The DCA is split into training and validation image sets. The AI image analysis depicting the architecture of the Inception V3 model is shown. The fully connected layer of the architecture is used for cell clustering and UMAP visualization. The softmax layer generates per cell classification and the prediction probabilities. It also outputs the cell z-plane focus metrics, which are used to report on image quality. The model prediction for debris, doublets and cell clumps is used to report on sample quality. **(D)** Realtime AI-based sorting workflow: Images of single cells are converted to a vector, and a user-selected classifier assesses each cell. The embedding vector generated by the model is used to visualize sample profile (e.g. UMAP depiction is drawn based on the embeddings). Additionally the realtime inferences guide a sorting decision, based on user preferences. The sorting decision then translates into valve control signals. The laser tracking system detects cells as they arrive in different outlets, through evaluating two photomultiplier tube (PMT) signals. The system generates reports of the number and type of analyzed cells, number of sorted cells, sorting purity and yield, focus plane, synchronization signals, and the fluidic pressures and flow rates. The system uses this information in a feedback loop to adjust system parameters.

## Hardware

A microfluidic cartridge allows for the input and flow of cells in suspension with confinement along a single lateral trajectory to obtain a narrow band of focus across the z-axis. Using a combination of hydrodynamic and inertial focusing, we collect high-speed bright-field images of cells (up to 20,000 frames per second) as they pass through the imaging zone of the microfluidic cartridge. Images capture subcellular and subnuclear features of the single cells in high contrast with each pixel representing an area of 0.044μm^2^. An automated object detection module tracks the cells as they flow through the channel. The images are fed into a CNN for generation of high-dimensional morphological descriptors and classification in realtime. Based on the classification, pneumatic valves are used for sorting a cell into either the cell collection reservoir or waste outlet (**Fig. 1, A and D and fig. S1**). Sorted cells are then retrieved for downstream analysis. A laser-based tracking system identifies cells in realtime, to assist with imaging, sorting and to report on the purity and yield of the run. The instrument can automatically align the microfluidic chip within the camera’s field of view, re-focus the optical z-plane, and adjust its operation based on sensors during instrument setup, imaging, and sorting.

## Cell Annotation

Images of single cells are the input to the AI-assisted image annotation software (**Fig. 1B**), which uses an unsupervised learning approach to assign annotations to images to train machine learning models. Agglomerative clustering is used to cluster cell images, which can be viewed grouped by their focal plane. These cell groups are generated in 2 modes: 1) clusters that are formed based on morphological similarities deduced by an expressive unsupervised model, and 2) morphological proximity to cells annotated within the same session or prior sessions. This software enables a human expert to re-assign annotations to cells that are incorrectly annotated or partition morphologically distinct clusters into multiple cell annotations. Trained users have achieved annotation rates over 100 cells per second using this tool.

## DCA

The DCA is an ever expanding database of expert-annotated images of single cells collected from a variety of immortalized cell lines, patient body fluids as well as tissue biopsies. At the time of this manuscript, DCA has amassed over 1.5 billion images of single cells. The annotations are structured based on a cell taxonomy which may allow a cell to be assigned multiple annotations on its lineage. The training pipeline extracts training and validation sets from DCA to train and evaluate neural net models aimed at identifying certain cell types and/or states. During training, one or more annotations may be selected for each cell image according to the architecture of the model (**Fig. 1C**).

## Machine learning

A machine learning infrastructure capable of realtime analysis of cell images was developed to generate high-dimensional morphologic descriptors and classifications (**Fig. 1C**). Our model architecture is based on the InceptionV3 (Szegedy et al. 2016) CNN, modified for grayscale images and to output quantitative morphological descriptors (often called “embeddings” in the machine learning literature). This architecture consists of 48 layers and 24 million parameters. Features from cell images are summarized as an embedding from which cell class annotations are predicted. These embedding vectors are not generally interpretable in terms of conventional morphology metrics but can be used to perform cluster analysis to group morphologically similar cells and visualized using tools like Uniform Manifold Approximation and Projection (UMAP) (McInnes et al. 2018), and clustered heatmaps. This architecture runs in realtime on our instrument which allows images to be analyzed by previously trained models and generates classification and high-dimensional morphology descriptions for each imaged cell. If cell sorting is desired, the model outputs are used to determine whether to discard or retain each cell and, if retained, which collection well to route each cell.

## Cells cluster in embedding space

To demonstrate that COSMOS can identify unique cell types, we applied it to cells that may circulate in body fluids, like fetal cells and cancer cells. Therefore, we created training and validation sets that included non-small cell lung cancer (NSCLC) cell lines, hepatocellular carcinoma (HCC) cell lines, fetal nucleated red blood cells (fnRBC), and adult peripheral blood mononuclear cells (PBMCs) (**Fig. 2B**), and measured the performance of COSMOS in identifying these different cell types (**Fig. 2, A and E)**. We generated low-dimensional projections of the embeddings from our trained model, using UMAP plots (**Fig. 2A**). We found a strong correlation between the dimensions of the embedding space and cell type, as illustrated using heatmap and UMAP representations (**Fig. 2, A and C**). The UMAP plot shows that distinct cell types are clustered separately from one another. Within the NSCLC and HCC cell line clusters, the three cell lines were clustered separately (e.g. A549, H522, H23). PBMCs show a large degree of variation consistent with being comprised of several morphologically distinct classes of cells. We then showed certain coordinates of the embedding space correlate with different cell classes by projecting the value of each coordinate in the embedding space onto the UMAP representation (**fig. S2**). Representative images of each of these four classes captured by COSMOS are shown in **Fig. 2D**.

**Figure 2.**
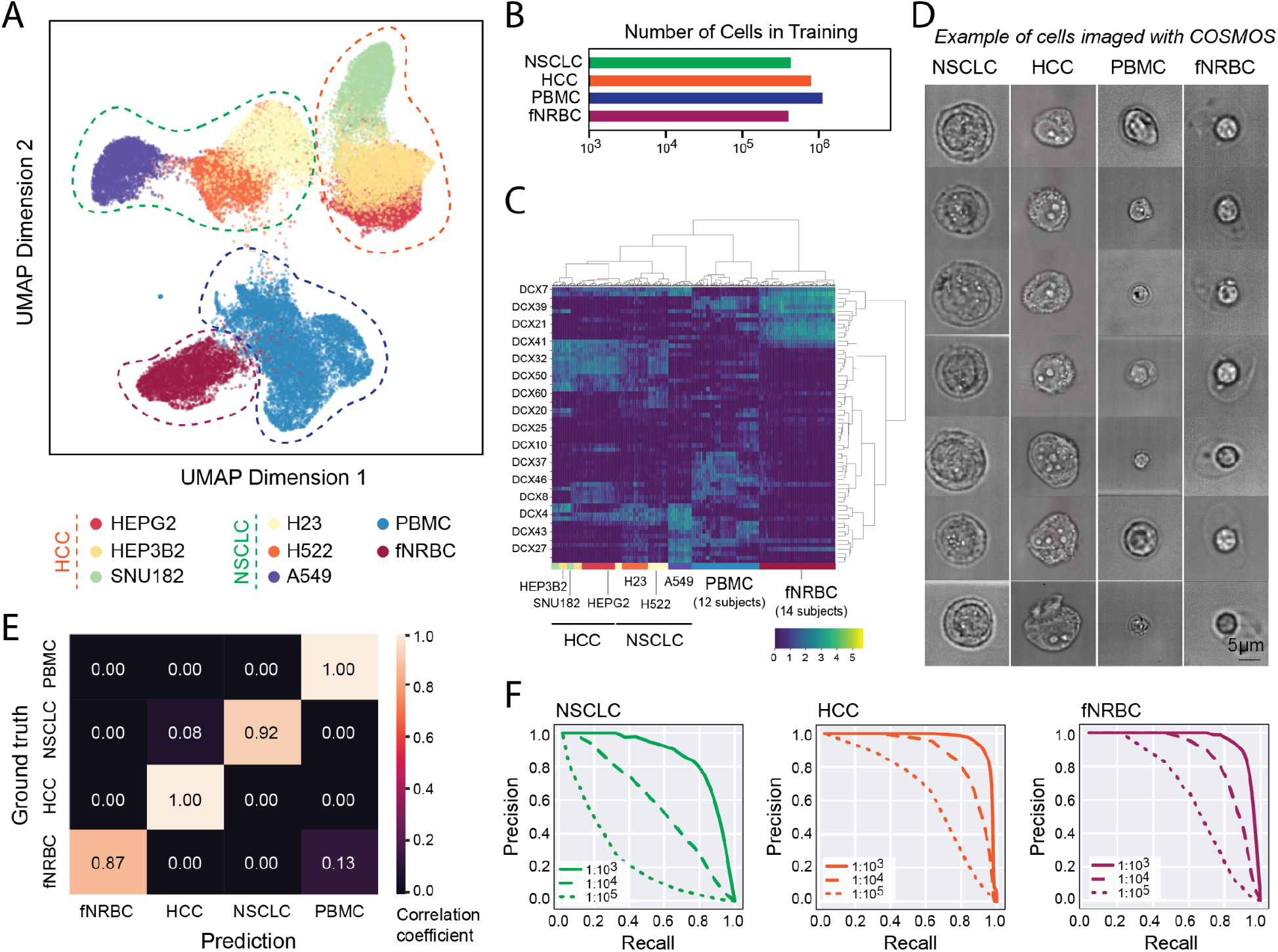
Quantitative morphological assessment of single cells, and performance of COSMOS in identifying cells. (**A**) UMAP projection of cell embeddings sampled from classes analyzed by the model. Each point represents a single cell. (**B**) The number of cells for each of the categories in the training set. (**C**) Heatmap representation of the embedding space. Each column is a single cell and each row is an embedding dimension. (**D**) Representative images of NSCLC, HCC, PBMC and fnRBC classes collected by COSMOS. (**E**) Confusion matrix representing the classifier’s prediction accuracy (x axis) versus ground truth (y axis). (**F**) Estimated precision-recall curves at different proportions for positive selection of NSCLCs, HCCs and fNRBCs against a background of healthy donor PBMCs. Precision corresponds to the estimated purity and recall to the yield of the target cells. Three curves are shown for different target cell proportions: 1:1,000, 1:10,000 and 1:100,000.

## Classification of cell types with low error

We next measured the accuracy of the model in classifying the four different cell types in a supervised fashion. For all the cell classes, the cell lines assessed in the validation dataset were distinct from those used for training. The validation dataset also included fnRBCs drawn from a pool of three fetal samples, and PBMCs extracted from the blood samples of three different donors, that were not used in the training dataset. **Fig. 2E** is the confusion matrix of classifier prediction correlations for each cell class against their true class. The data shows that the model’s prediction for fNRBCs, HCCs, NSCLCs and PBMCs matches the actual class at 87%, 100%, 92% and 100%, respectively. The confusion matrix demonstrates that morphology alone can accurately differentiate and identify these cell types when compared against each other.

## *In silico* evaluation of cell enrichment in contrived blood samples

We assessed the ability of COSMOS to identify low abundance NSCLCs, HCCs and fnRBCs from a background of PBMCs. We considered two different strategies for evaluating performance of the supervised model: positive (selecting the target cell class: NSCLC or HCC) and negative selection (selecting all nucleated blood cells: PBMC). The classifier performance metrics for these cell lines yielded an area under curve (AUC) of 0.9842 for positive selection and 0.9996 for negative selection, respectively, for the NSCLC class, and an AUC of 0.9986 and 0.9999 for positive and negative selection, respectively, for the HCC class (**fig. S3, A and B**). In addition, we demonstrated low false positive rates for both modes of classification. Although the AUCs are superior in the negative selection strategy in both cases, the positive selection strategy in both cases would enable higher yields at low false positive rates (FPR < 0.0004). For fnRBCs, we assessed only the mode of positive selection which yielded an AUC of 0.97 (**fig. S3C**).

To better understand the model performance, different spike-in ratios were analyzed *in silico*. Estimated precision-recall curves at different proportions of target cells (NSCLC, HCC and fNRBCs) in a background of healthy donor PBMCs demonstrates that even at a dilution of 1:100,000, the model supports detection of target cells at >70% precision (positive predictive value or post-enrichment purity) and 50% recall (sensitivity) in both the fnRBC and HCC samples, while precision drops to 15% for NSCLC class (**Fig. 2F**). We also show the probability distribution for each of the classes as it relates to their identification against PBMCs for both positive selection (P_NSCLC_, P_HCC_ and P_fnRBC_) and negative selection (P_PBMC_) (**fig. S4**).

## Enrichment of target cells

To biologically validate our i*n silico* analysis, we performed simultaneous classification and enrichment experiments by spiking NSCLC cell lines (A549 and H522) into PBMCs at defined proportions ranging from 1:1,000 to 1:100,000. The fnRBC sample was spiked into PBMCs from matching maternal blood. Each spike-in mixture was then processed on COSMOS and cells identified as target cells (fnRBC or NSCLC) by the classifier were sorted in realtime and subsequently retrieved.

For each spike-in mixture, we assessed the purity of the sorted cells retrieved from our system by analyzing allele fractions of the spiked-in cell lines and the background cells in a panel of single nucleotide polymorphism (SNP) assays (**fig. S5)**. By comparing the known spike-in mixture proportions and the final purity, we computed the degree of enrichment achieved for each of the samples analyzed. COSMOS was able to achieve similar enrichment and purity for A549 and H522 cells (**Fig. 3A**, **table S1**), even though the former was used to train the classifier and the latter was not. For the lowest spike-in ratio investigated (1:100,000), 20% and 30-33% purities corresponding to folds enrichments of 13,904x and 30,000x-32,500x were obtained for A549 and H522, respectively.

**Figure 3.**
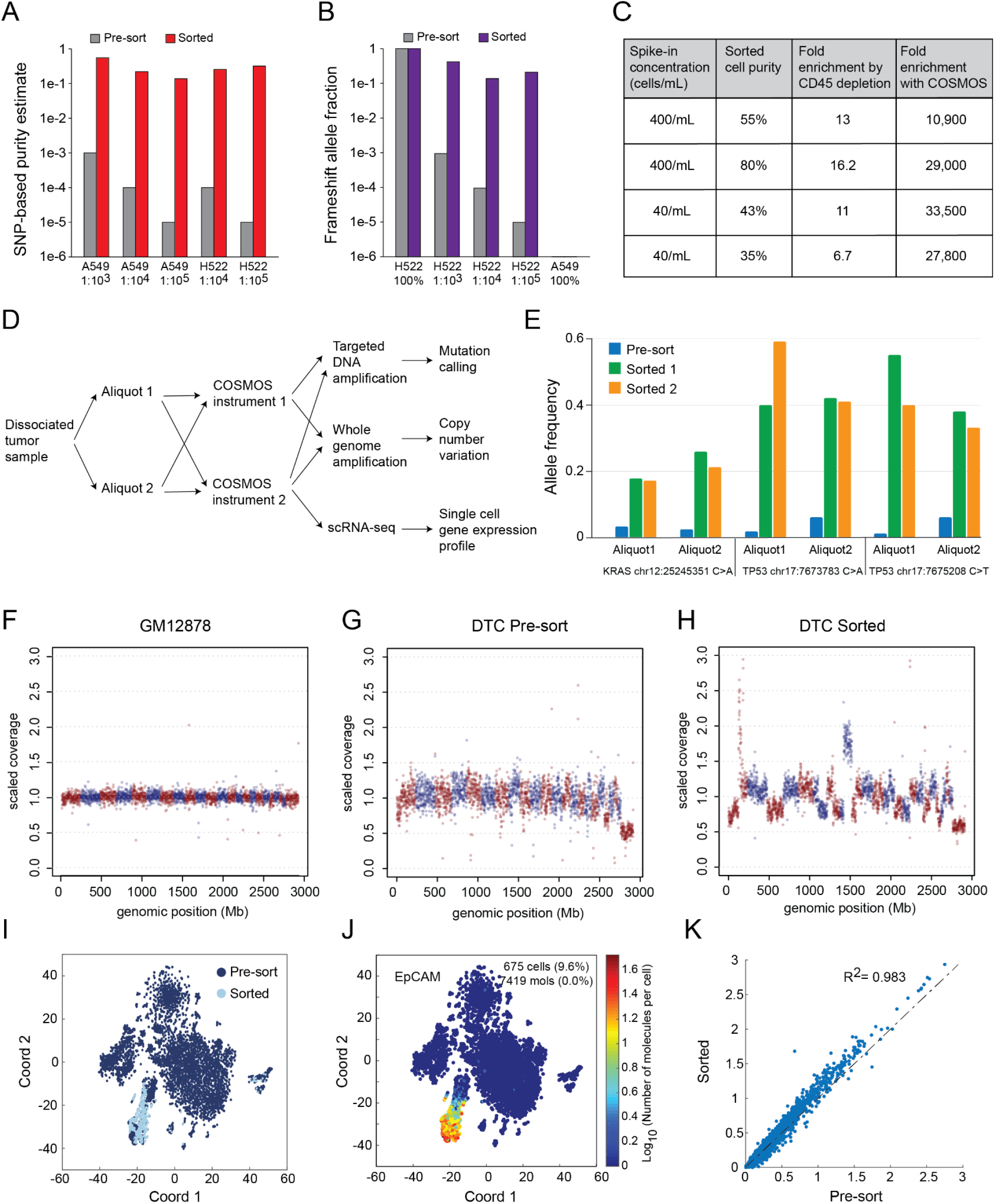
Performance of COSMOS in identifying and isolating target cells. A549 and H522 cell lines were spiked into donor PBMCs at the indicated ratios and processed on COSMOS for target cell identification and sorting. Purity of pre-sorted and sorted cells was estimated by comparing (**A**) allele fractions with a SNP panel to the known genotypes of both the cell lines and the donor samples that they were spiked into and (**B**) a frame-shift mutation assay in the *TP53* gene (c.572_572delC), for which the H522 cell line is homozygous and the A549 cell line is wild type. (**C**) The indicated number of A549 cells were spiked into whole blood. CD45 depletion was performed and samples were processed on COSMOS for malignant cell identification and sorting. Purity of the sorted cells and fold enrichment were quantified by SNP analysis using known genotypes of both the A549 cell line and the blood samples that they were spiked into. (**D**) Workflow schematic of COSMOS sorting and downstream molecular analysis of DTCs. (**E**) A KRAS mutation (Chr12:25245351 C>A) and two TP53 mutations (chr17:7673783 C>A and chr17:7675208 C>T) were discovered in this sample and the allele frequency in pre-sorted and sorted samples is shown between sorting runs and aliquots. (**F-H**) WGA and CNV analysis of the pre-and post-sorted samples. Each data point represents 1Mb bin. Red and blue colors indicate different chromosomes. GM12878 genomic DNA was used as baseline control for copy number normalization. (**I**) scRNAseq was performed and a t-SNE plots of gene expression profiles using all 924 feature-selected genes for pre-sorted (dark blue) and post-sorted (light blue) is shown as an overlay and (**J**) the pseudo-color gene expression level of EPCAM (cancer cell marker) is shown. (**K**) Gene expression correlation plot of mean (log_10_(molecules per cell per gene)) for the sorted and the pre-sorted cells from the EPCAM^+^/PTPRC(CD45)^-^ cluster. Each data point is a gene. The gene expression correlation coefficient (R^2^) was 0.98.

We also assayed for a frameshift mutation in *TP53* (c.572_572delC), for which the H522 cell line is homozygous and the A549 cell line is wildtype (Tate et al. 2019). The proportion of the total sequence reads that contain this frameshift mutation are shown in **Fig. 3B** and **table S2** and are consistent with purity estimates from the panel of SNPs depicted in **table S1**. Even at the lowest investigated spike-in ratio of 1: 100,000, we found the mutation present at an allele fraction of 23% in the DNA extracted from the enriched cells, suggesting that functionally important cancer mutations may be detected even when the cells containing them are present at proportions significantly lower than 1:100,000.

Next, we spiked A549 cells into whole blood at concentrations of 40 cells/mL and 400 cells/mL and processed them as outlined in the methods. The purity and fold enrichment of the sorted cells was estimated by jointly analyzing allele fractions in a SNP panel for both the A549 cell line and the enriched cells (**Fig. 3C** and **fig. S6**). The sorted samples had final purities of 55% and 80% for the 400 cells/mL replicates (corresponding to an overall enrichment of >10,900 fold and >29,000 fold respectively) and purities of 43% and 35% for the 40 cells/mL replicates (corresponding to an overall enrichment of >33,500 fold and >27,800 fold, respectively) (**Fig. 3C** and **table S3**).

## Compatibility of sorted cells with single cell RNA sequencing (scRNAseq)

We tested if the cells sorted with COSMOS were viable and amenable to downstream scRNAseq analysis. We found COSMOS had minimal or no impact on cell viability across the cell lines and primary cells tested (**table S4**). We further compared the single cell gene expression profiles between unprocessed and COSMOS processed PBMCs by scRNAseq with a targeted immune response panel and whole transcriptome amplification (WTA). We found high correlation between the gene expression profiles using both targeted assays (R^2^=0.97) and whole transcriptome (R^2^=0.983), indicating that the cells processed through COSMOS are directly comparable with unprocessed cells, and are compatible with downstream single cell RNA analyses (**fig. S7, A, B and C**). Additionally, we compared the cell health states of neutrophils, a cell type that is known to be sensitive to cell processing (Alvarez-Larran et al. 2005), after various processing workflows (**fig. S7D**). With bulk RNA sequencing analysis, COSMOS-sorted cells showed fewer up- or down-regulated genes relative to control cells (**fig. S7E**) compared to FACS. They had less activation in genes involved in multiple immune cell activation pathways and neutrophil degranulation pathways (**fig. S7F**), suggesting COSMOS sorting was gentler to the cells.

## Identification of malignant cells from dissociated solid tissue biopsies

We evaluated the accuracy of the model in identifying malignant cells from dissociated solid tissue biopsies, by running the model on three NSCLC dissociated tumor cell samples (DTC) with low, medium, and high percentages of malignant cells and comparing the model results to flow cytometry and scRNAseq analysis (**fig. S8**). Malignant cell frequencies determined by the model had high concordance to scRNAseq analysis of EpCAM expression for low (2.2% vs 4.6%), medium (12% vs 16.8%) and high (40% vs 46.7%) malignant cell purities (**fig. S8)**.

## Enrichment of malignant cells from DTC samples

Finally, as proof that COSMOS can specifically distinguish and enrich malignant cells from tumor tissue, we sorted cancer cells from a DTC sample of a stage IIB NSCLC patient. To confirm the run-to-run consistency, the sample was split into two aliquots, and each aliquot was run on two COSMOS instruments. Sorted cells were split into multiple fractions for molecular analysis, including targeted DNA panel amplification for mutation analysis, whole genome amplification (WGA) followed by copy number variation (CNV) analysis, and scRNAseq for gene expression analysis (**Fig. 3D**). Our model predicted 1.2% ± 0.7% of malignant cell fraction on multiple runs, consistent with the EpCAM+ percentage reported by FACS (1.3%-1.5%; data not shown). Using a targeted lung cancer panel we found one KRAS and two different TP53 mutations and in sorted samples the allele frequency increased from <3% to 20% and 1-6% to 33-59%, respectively **(Fig. 3E)**. The two pre-sorted aliquots showed variations in the allele frequencies, possibly due to both tumor cell heterogeneity and technical noise in amplification of rare cells at 1-2% range. Nonetheless, we were able to enrich the mutations to 20-60%, suggesting COSMOS enrichment both captured the mutational heterogeneity of the pre-sorted sample and improved confidence in mutation calling beyond any technical noises for low tumor content samples. We then profiled the bulk copy numbers by WGA and were able to significantly increase the sensitivity of CNV detection (**Fig. 3, F, G and H)**. For example, chr8q was amplified (**Fig. 3H**), upon which the *MYC* and *PRDM14* oncogenes are located (Baykara et al. 2015).

We confirmed the identities of the sorted cells, their suitability for single cell gene expression analysis and compared the scRNA profiles to the pre-sorted sample using a WTA workflow. We found that 86-92% of the sorted cells overlapped with EpCAM+/CD45-populations from the pre-sorted cells, indicating a high degree of purity in the sorting capability **(Fig. 3,I and J and fig. S9a)**. The sorted and pre-sorted cells from the EpCAM+/CD45-cluster showed strong gene expression correlation (R^2^ = 0.98), and overlapped in all subclusters, suggesting that COSMOS sorting was unbiased at least for the EpCAM+ population and did not change gene expression profile due to the gentle microfluidic flow (**Fig. 3K and fig. S9, B and C**). A close examination of 166 stress and apoptosis-related genes (a preloaded gene set from DataView software) also did not show differences in the sorted cells compared to the pre-sorted sample (**fig. S9, E, F and G**).

In conclusion, we present COSMOS, a novel technology platform for the characterization, classification, isolation, and enrichment of cells from living organisms based on high-dimensional morphology. Recent work has motivated morphology as an analyte in cell sorting (Schraivogel et al. 2022). Here we capture the power of deep neural networks in processing morphology by amassing an annotated atlas of greater than 1.5 billion single cell images and training deep models with the computational capacity to classify high resolution high content images. COSMOS offers deep interpretation of single cell phenotype in realtime, with no need for complex sample pre-processing, gating, feature engineering, or bioinformatics capabilities. Using its label-free unbiased approach, COSMOS provides a unique capability to analyze and enable discovery in cell populations with unknown phenotypic or molecular makeup. By enriching viable unaltered cells from tissue and the circulation, the platform enables the combination of morphologic and molecular characterization at the single cell scale, providing novel insights to advance our understanding of biology in basic, translational and clinical applications.

## Methods and Materials

### Microfluidics

Each cartridge design has a microfluidic channel height between 15 μm and 40 μm, chosen to be a few micrometers greater than the largest cells to be processed. A filter region at the input port prevents large particles, cells or cell aggregates from entering the flow channel. A buffer reagent (1X PBS) is introduced into the flow alongside the cell suspension on either side, achieving hydrodynamic focusing that keeps cells flowing at a consistent speed near the center of the flow horizontally. The flow rate used (~0.1 m/s) is also high enough that the effects of inertial focusing (Di Carlo et al. 2007) are realized, confining cells to the vicinity of two vertically separated planes close to the center of the flow channel.

### Bright-field imaging of cells in flow

The microfluidic cartridge is mounted on a stage with lateral (horizontal) XY control and a fine Z control for focus. The objectives, camera, laser optics and fluidics components are all mounted on the same platform. After the microfluidic cartridge is loaded into COSMOS, it is automatically aligned and a focusing algorithm is used to bring the imaging region into the field of view. An LED illumination light (SOLA SE) is directed to the imaging region, and multiple images of each cell are captured as it flows through. Bright-field images are taken through objectives of high magnification (Leica 40X - 100X) and projected onto an ultra high-speed camera. To achieve higher accuracies and adjust for potential artifacts in the image, at least two images are captured from each cell as they flow downstream in the channel. These high-resolution cell images reveal not only the cell shape and size but also finer cellular structural features within the cytoplasm and the nucleus that are useful for discriminating cell types and states based on their morphology.

### Computation

The COSMOS software workload is distributed over an Intel Xeon E-2146G central processing unit (CPU), a Xeon 4108 CPU, an Nvidia Quadro P2000 Graphical Processing Unit (GPU) and a custom microcontroller. The camera is periodically polled for the availability of new images. Image frames from the high speed bright-field camera are retrieved over a dedicated 1Gbps ethernet connection. Images are cropped to center cells within them, and the cropped images are sent to the GPU for classification by an optimized CNN that has been trained on relevant cell categories. The network architecture is based on the Inception V3 model architecture (Szegedy et al. 2016), is implemented using the TensorFlow v1.15 (Abadi et al. 2016), and is trained using cell images annotated with their corresponding cell categories. NVidia TensorRT is used to create an optimized model which is used for inference on the GPU. The classification inference from the models is sent to the microcontroller, which in turn sends switching signals to synchronize the toggling of valves with the arrival of the cell at the sorting location. To maximize throughput, image processing happens in a parallel pipeline such that multiple cells can be in different stages of the pipeline at the same time. The primary use of the GPU is to run the optimized CNN. Some basic image processing tasks such as cropping cells from the images are performed on the instrument CPU. The instrument CPU is also used to control all the hardware components and to read in sensor data for monitoring. The training and validation tasks are set up as recurring Apache Beam based data processing pipelines in Google Cloud Platform (GCP). Training and prediction jobs are orchestrated by Apache Airflow, and Google Cloud Dataflow is used to combine predictions, embeddings and annotations. Models are trained using TPUPodOperators on Google Cloud on version 3 of Google’s Tensor Processing Units. PostgreSQL, Google Big Query, and Google Cloud Storage are used to store and query model predictions, embeddings, and run metadata.

### Data augmentation and model training

Several steps were taken to make the image classifier robust to imaging artifacts by systematically incorporating variation in cell image characteristics into our training data. Cells were imaged under a range of focus conditions to sample the effects of changes in focus during instrument runs. We gathered images across four of our instruments to sample instrument-to-instrument variation. We also implemented several augmentation methods to generate altered replicas of the cell images used to train our classifier. These included standard augmentation techniques such as horizontal and vertical flips of images, orthogonal rotation, gaussian noise, and contrast variation. We also added salt-and-pepper noise to images to mimic microscopic particles and pixel-level aberrations. Finally, we studied systematic variation in our image characteristics to develop custom augmentation algorithms that simulate chip variability and sample-correlated imaging artifacts on our microfluidic cartridge.

All cell images were resized to 299×299 pixels to make them compatible with the Inception architecture. We trained a model comprising cell types present in normal adult blood, cell types specific to fetal blood, trophoblast cell lines, and multiple cancer cell lines drawn from NSCLC, HCC, pancreatic carcinoma, acute lymphoblastic leukemia (ALL), AML. The model was also trained to detect out-of-focus images, both to use this information in auto-focusing during instrument runs and to exclude out-of-focus cell images from possible misclassification.

### AI-assisted annotation of cell images

For the supervised model, we collected high-resolution images from 25.7 million cells, including cells from normal adult blood, fetal blood, trophoblast cell lines, and multiple cell lines derived from NSCLC, HCC, pancreatic carcinoma, ALL, and AML. Images were collected by an ultra high-speed bright-field camera as cell suspensions flowed through a narrow, straight channel in a microfluidics cartridge. We deployed a combination of techniques in self-supervised, unsupervised, and semi-supervised learning to facilitate cell annotation on this scale. First, we used subject and sample source data to restrict the set of class labels permitted for each cell; as an example, fetal cell class annotations were disallowed in cells drawn from non-pregnant adult subjects. Next, we extracted embedding vectors for each cell image in two pre-trained CNNs: one trained on the ImageNet dataset (Russakovsky et al. 2015) and the other on a subset of our own manually annotated cell images. We then used agglomerative clustering of these feature vectors to divide the dataset into morphologically similar clusters which were presented for manual annotation, thereby facilitating efficient cell annotation at scale.

To further enhance the accuracy of subsequent cell classification, we also selectively annotated false positive images identified from the predictions of previous trained models in an iterative manner. Finally, we balanced the classes that we wish to discriminate by feeding the harder examples of more abundant classes inspired by an active learning approach. The hard examples were identified as those that a model trained on a smaller training set had classified incorrectly (Settles 2010).

### Training and validation sets

57.4 million images were gathered to train and validate the classifier. A dataset of 25.7 million cells was imaged for the purpose of training our deep CNN in the model: PBMCs of 44 blood samples of normal adult individuals were collected which resulted in 22 million cell images. Additionally, 18 fetal blood samples were collected which yielded 2.8 million imaged cells. We imaged a total of 156,000 cells from four NSCLC cell lines, a total of 400,000 cells from four HCC cell lines, and another 440,000 cells from four cell lines of other types. A separate dataset of 25.1 million cells from 111 samples of the cell types above were gathered to validate the results of the classifier. We used the NCI-H522 (H522) cell line as the sample in validation for NSCLC and Hep 3B2.1-7 (HEP3B2) for HCC respectively.

### Cell sorting

Cell sorting is performed using built-in pneumatic microvalves (Unger et al. 2000) on both the positive (targeted) and negative (waste) sides of the flow channel downstream of the bifurcation point. Valve timing is controlled by a DSP-based microcontroller circuit with 0.1ms time precision. When the model infers that a cell belongs to a targeted category, switching signals are timed to synchronize the toggling of valves with the arrival of the cell at the flow bifurcation point, and the cell flows into a reservoir on the microfluidic cartridge where targeted cells are collected (also called the positive well). If the model infers that a cell does not belong to a targeted category, the cell flows into a waste tube. Elliptical laser beams are focused onto both the positive and negative output channels downstream of the sorting flow bifurcation to detect passing cells and thereby monitor sorting performance in realtime.

### Sample processing and cell culture

All human blood samples were collected at external sites according to individual institutional review board (IRB) approved protocols and informed consent was obtained for each case. For adult control and maternal blood samples, white blood cells (PBMCs) were isolated from whole blood by first centrifugation then the buffy coat was lysed with Red Blood Cell (RBC) Lysis Buffer (Roche) and then washed with PBS (Thermo Fisher Scientific). Fetal cells were isolated from fetal blood by directly lysing with the RBC lysis buffer then washed with PBS. Cells were then fixed with 4% paraformaldehyde (Electron Microscopy Sciences) and stored in PBS at 4°C for longer term usage. A549, NCI-H1975, NCI-H23 (H23), NCI-H522 (H522), NCI-H810, Hep G2 (HEPG2), SNU-182, SNU-449, SNU-387, Hep 3B2.1-7 (HEP3B2), BxPC-3, PANC-1, Kasumi-1, Reh, and HTR-8/SVneo cell lines were purchased from ATCC and cultured in a humidity and CO_2_-controlled 37°C cell culture incubator according to ATCC recommended protocols. GM12878 cell line was obtained from the NIGMS Human Genetic Cell Repository at the Coriell Institute for Medical Research and cultured according to their recommended protocols.

For neutrophil isolation and sorting, human neutrophils were isolated from whole blood using the EasySep Direct Human Neutrophil Isolation kit from Stemcell Technologies by immunomagnetic negative selection. When applicable, isolated neutrophils were labeled with a panel of primary antibodies (anti-CD3, anti-CD45, anti-CD19, anti-CD14, anti-CD66b, anti-CD15 from Biolegend) for 20 minutes at room temperature and washed twice. Propidium iodine was added to the cell mixture prior to acquisition and sorting on a BD FACSMelody instrument.

For spike-in experiments, cancer cell lines or fetal cells were first fixed with 4% paraformaldehyde and stored at 4°C until mixing into PBMCs. For experiments in which cell lines were spiked into whole blood, live A549 cells were first stained with CellTracker Green CMFDA (Thermo Fisher Scientific), then spiked into whole blood (collected in EDTA tubes) at predefined ratios (e.g. 400 or 4000 cells in 10 mL blood), followed by buffy coat RBC lysis and fixation. Prior to loading into the sorter, the cell mixtures were pre-enriched by selective depletion of CD45 positive PBMC cells using magnetic beads (Miltenyi). Twenty percent of the samples were saved for flow cytometry analysis to estimate the number of total cells and cancer cells before and after CD45 depletion. Based on flow cytometry analysis, the CD45 magnetic bead depletion step resulted in 11-15 fold enrichment of A549 cells.

DTCs from NSCLC patients were purchased from Discovery Life Sciences. Cancer type and stage information and cell type composition report from flow cytometry were provided by the vendor. To account for possible cell type composition changes from the freeze-thaw process, after thawing the DTC aliquots, we split the samples to analyze some cells with flow cytometry and image and sort some cells on COSMOS. The panel used for flow cytometry includes markers: EpCAM, CD45, CD3, CD16, CD19. CD14, CD11b.

For cell viability assessment, pre-sorted or sorted cells were stained with either trypan blue or a Calbiochem live/dead double staining kit (Millepore Sigma) which uses a cell permeable green fluorescent Cyto-dye to stain live cells and propidium iodine to stain dead cells. Cells were then counted under a fluorescent microscope.

#### Molecular analyses

##### Single cell RNA sequencing

Cells were either directly loaded or retrieved from the positive wells of the microfluidic cartridge then loaded on a BD Rhapsody single cell analysis system (BD Biosciences). Single cells were then processed following either targeted RNA sequencing (human immune response panel) or whole transcriptome amplification protocols. The sequencing data were analyzed using BD DataView software.

##### Bulk RNA sequencing

Total RNA was extracted from cells using the RNeasy mini kit from Qiagen. cDNA synthesis, amplification and library preparation were performed with the Quantseq 3’m RNAseq library prep kit from Lexogen according to the manufacturer’s protocol. The final libraries were sequenced on an Illumina Miniseq. Read QC, trimming, alignment and counting were performed with the Lexogen Quantseq analysis pipeline. Differential expression analysis was done using DESeq2 and iDEP (http://bioinformatics.sdstate.edu/idep/).

##### Genotyping

Cell lines and PBMCs of individual blood donors were genotyped with Next Generation Sequencing using a targeted SampleID panel (Swift Biosciences) that includes 95 assays for exonic single nucleotide polymorphisms (SNPs) and 9 assays for gender ID. Briefly, genomic DNA was extracted from bulk cells using QIAGEN DNeasy Blood & Tissue Kit (Qiagen) and 1ng DNA was used as input to amplify the amplicon panels and prepare the sequencing library. For cancer cell lines, a 20-amplicon panel that covers full length of TP53 gene (Swift Biosciences) was pooled with the SampleID panel so cells were genotyped on both common SNPs and TP53 mutational status. From ATCC and COSMIC annotation, A549 cells are known to be TP53 wild type and NCI-H522 are known to carry a homozygous frameshift mutation (c.572_572delC). Our bulk genotyping results confirmed the relative mutation status for these two cell lines. For sorted cells from the COSMOS experiments, cells were retrieved from the positive outlet well of the microfluidic cartridge into a PCR tube, then directly lysed using Extracta DNA Prep for PCR (Quanta Bio). Cell lysates were amplified with the Swift amplicon panels and followed by the same library preparation procedure for NGS.

##### Dissociated tumor cells from lung cancer patients

The cells before sorting and after sorting were profiled on targeted DNA mutations and copy number variations (CNV). For mutation analysis following direct lysis with Extracta DNA Prep for PCR (Quanta Bio) a 208-amplicon panel that includes 17 lung cancer genes (Swift Biosciences) were used. For CNV analysis, after direct lysis, genomic DNA was amplified using ResolveDNA Whole Genome Amplification Kit (BioSkryb Genomics) and then libraries were prepared for sequencing (Kapa Hyperplus Kit, Roche). All libraries were sequenced on either an Illumina MiniSeq or NextSeq instrument (Illumina) using 2×150 bp kit (DNA) or 2×75 bp kit (RNA).

##### Primary sequencing analysis and QC

Sequencing reads were aligned to the reference genome using the BWA-MEM aligner. SNP allele counts were summarized using bcftools. SNP data were subjected to quality control checks: each sample was required to have a mean coverage per SNP of > 200; each SNP locus needed to have a median coverage across all samples > 0.1x the median SNP to be considered; each individual SNP assay for a sample needed to have a depth of coverage > 50. 89 SNP assays were selected on this basis for further use in mixture analysis. Samples and individual SNP assays that failed QC were excluded from genotyping and the estimation of mixture proportions.

##### Mixture proportion estimation by SNP analysis

Pure diploid samples that formed the base of each mixture for spike-in experiments were clustered into the three diploid genotypes (AA, AB, BB) for each SNP using a maximum likelihood estimation that incorporated an internal estimate of error within homozygous SN. The mixture proportion of the component of interest (tumor cell line or fetal sample) was determined using maximum likelihood estimation (MLE), in which all discrete mixture fractions in increments of 0.005 were considered (0.0, 0.005, 0.01,…, 1.0). For each possible mixture proportion, expected allele fractions at each SNP were determined by linearly combining the allele fractions in the two mixture components. A binomial log likelihood corresponding to each individual sample-SNP combination was computed using the expected allele fraction and an effective number of independent reads N per SNP estimated from the variance of allele fraction in mixture SNPs at which the base genotype is heterozygous (AB) and the spike-in component genotype is homozygous (AA or BB). By estimating N from the mixture data directly and using SNPs expected to have a shared allele fraction, the procedure is robust to low input for which the number of reads might exceed the number of independent molecules sampled. The overall log likelihood for each possible mixture proportion is computed as the sum of contributions from each SNP, and the mixture proportion is estimated as that at which the highest overall log likelihood is obtained. The accuracy of the procedure was verified on DNA mixtures with known composition (**fig. S5**). Each composite sample contained 250 pg of DNA and the mixture proportion of DNA from the second individual was set at 5%, 10%, 20%, 30%, 40%, 60%, 80% and 90%.

##### Joint Estimation of Genotypes and Sample Purity

In two cases, genotypes and mixture fraction were jointly estimated from the allele fractions ϕ of SNPs in the mixture: (i) to genotype the fetal sample Fet1, which included some maternal cells in addition to fetal cells (ii) for the spike-in of A549 cells into whole blood. In each case, genotypes for one of the mixture components, designated ***G_0_***, were obtained from a pure sample (from maternal DNA for the former, and from the pure A549 cell line for the latter), while the genotypes of the other sample, designated ***G*** (corresponding to the fetal sample in the former case and to the unrelated blood sample for the latter) were estimated from the data. The maternal sample was genotyped as diploid, but for pure A549, the allowed allele fractions for genotypes were 0, ⅓, ½, ¾ and 1, in keeping with the known hypotriploidy of that cell line. An expectation maximization (EM) procedure was then used to jointly estimate the purity and missing genotypes. Briefly, given ***G_0_*** and a current estimate of purity *f*, a binomial likelihood was estimated for each allowed missing genotype, and a maximum likelihood estimate was used to update ***G***. Given ***G***, a revised estimate of *f* was obtained by linear regression, using the expected linear relationship between the observed allele fraction ϕ and ***G_0_*** over SNPs of identical ***G***. The procedure incorporated an error rate estimate drawn from the SNPs where both components are identically homozygous. The procedure was iterated until convergence, defined as changes in the purity estimate < 0.0001. Results of the EM procedure for A549 cells enriched from a starting concentration of 40 cells/mL are shown in **fig. S6**. The three dotted lines depict the linear regression used to estimate the purity given the genotypes; their slope is equal to the final purity estimate of 0.43. **Fig. 6D** also shows well-separated clusters corresponding to each of the inferred genotypes in the blood sample.

##### Mutation and CNV analysis

Mutation allele fractions in sorted and enriched samples (**fig. 3, B and E**) were estimated from targeted amplicon sequencing data. In each sample, the mutation allele fraction was estimated as the fraction of high-quality read alignments overlapping the mutation locus that contained the variant allele. For spike-in samples at concentrations < 1 % (1:1000, 1:10,000 or 1:100,000 in **fig. 3I**), the depicted pre-enrichment allele fraction is the experimental spike-in fraction.

Six aliquots from the GM12878 cell line, consisting of 100, 50, 25, 10, 5 and 1 cell(s) respectively, were used as a normalization cohort for copy number estimation in dissociated tumor cells before and after enrichment by sorting. Read coverage was first aggregated over 1 Mb genomic intervals across the genome within the dissociated tumor sample and each of the GM12878 normalization aliquots. The coverage within each sample was then scaled by the mean coverage per Mb over the entire genome for that sample. Next, the median assay bias and median absolute deviation (MAD) of the scaled coverage for each 1 Mb interval across the genome were computed from data from the normalization cohort. Genomic intervals for which the MAD across normalization samples exceeded 20% of the median were excluded from further analysis. Finally, the coverage values within the dissociated tumor sample before and after enrichment were further scaled by the median assay bias estimated from the normalization cohort. The resulting scaled coverage data reveal several large-scale aneuploidies in the dissociated tumor cells after sorting but not prior to the sort (**fig. 5, C, D and E**), and thereby provide strong evidence for an enrichment of tumor cells by sorting.

## Supplementary Data and Figures

**Table S1.** Enrichment of cells spiked into PBMCs. Fet1 is a fetal blood sample spiked into cells from the corresponding maternal sample. Cells from the A549 and H522 cell lines were spiked into PBMCs from a healthy donor. Cell mixtures were flown through, imaged and target cell sorted via COSMOS system. In some experiments, actual spike-in ratios of the mixtures were estimated and confirmed by pre-staining cancer cells with a fluorescent Cell Tracker dye before mixing into PBMC or whole blood and then analyzing a portion of cells using flow cytometry. Classifier Positive Rate is the percentage of the classifier identified positive cells against all imaged cells. Sorted Cell purity was estimated by comparing allele fractions using a SNP panel to the known genotypes of both the cell lines and the samples that they were spiked into and normalized by copy numbers of the cell lines. Fold enrichment was calculated by the SNP-estimated purity in sorted cells divided by target purity or flow cytometry-based estimation of pre-sort cells.

| Cell Source | Primary Cell Class | Target Spike-in Ratio | Cells Imaged | Classifier Positive Rate | Sorted Cell Purity | Fold Enrichment |
| --- | --- | --- | --- | --- | --- | --- |
| Fet1 | fnRBC | 1:1,304 | 999,978 | 0.017% | 74% | 965 |
| A549 | NSCLC | 1:1,000 | 69,611 | 0.150% | 62% | 348 |
| A549 | NSCLC | 1:1,000 | 101,180 | 0.170% | 67% | 380 |
| A549 | NSCLC | 1:10,000 | 1,105,997 | 0.060% | 27% | 1,978 |
| A549 | NSCLC | 1:10,000 | 876,421 | 0.099% | 17% | 1,201 |
| A549 | NSCLC | 1:10,000 | 1,107,669 | 0.025% | 31% | 2,305 |
| A549 | NSCLC | 1:10,000 | 1,063,745 | 0.0083% | 42% | 4,200 |
| A549 | NSCLC | 1:10,000 | 1,169,744 | 0.0094% | 33% | 3,300 |
| A549 | NSCLC | 1:10,000 | 719,499 | 0.028% | 33% | 3,300 |
| H522 | NSCLC | 1:10,000 | 1,050,036 | 0.030% | 26% | 2,550 |
| A549 | NSCLC | 1:100,000 | 1,342,632 | 0.003% | 20% | 13,904 |
| H522 | NSCLC | 1:100,000 | 1,514,263 | 0.005% | 30% | 30,000 |
| H522 | NSCLC | 1:100,000 | 1,561,847 | 0.006% | 33% | 32,500 |

**Table S2.** Detection and enrichment of a known frame-shift mutation in the TP53 gene for which the NCI-H522 cell line is homozygous. The indicated cell lines were spiked into healthy donor PBMCs 0.1% (1:1,000), 0.01% (1:10,000) and 0.001% (1:100,000). Each of these mixtures was then enriched using COSMOS. DNA from the enriched cells was assayed for the frame-shift mutation. In each case, the mutation was detected with an allele fraction of 15% or more. For the 1:100,000 spike-in mixture, an enrichment of 22,770x was achieved.

| Cell Line | Proportion in Mixture | Proportion of TP53 frameshift c.572_572delC | Fold enrichment of c.572_572delC |
| --- | --- | --- | --- |
| A549 | 100% | 0% | -- |
| NCI-H522 | 100% | 100% | -- |
| NCI-H522 | 0.1% (1:1,000) | 45% | 451 |
| NCI-H522 | 0.01% (1:10,000) | 15% | 1,499 |
| NCI-H522 | 0.001% (1:100,000) | 23% | 22,770 |

**Table S3.**
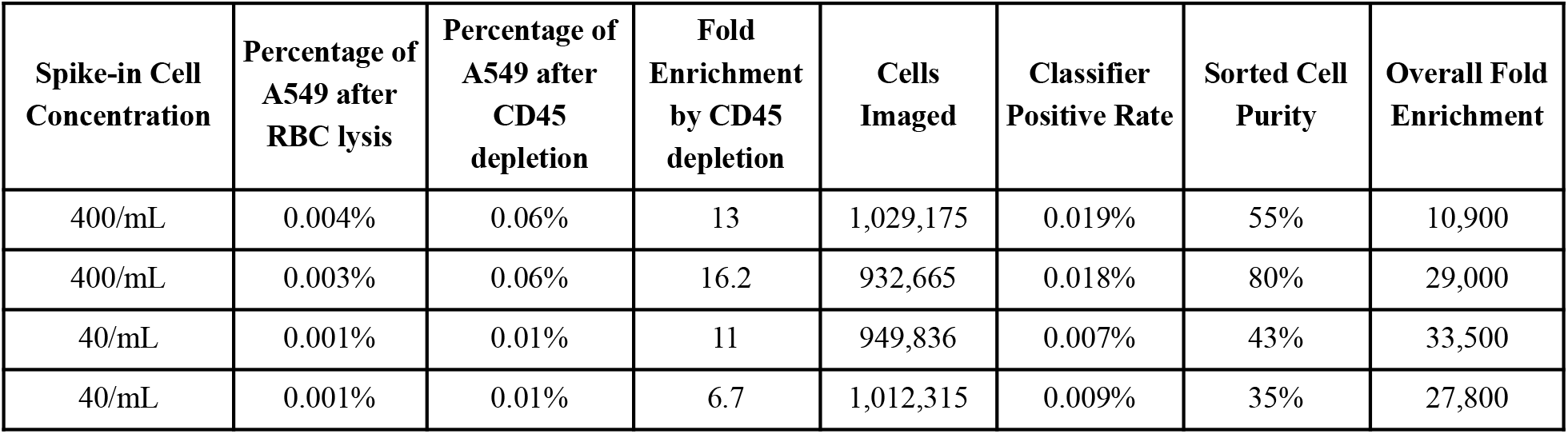
Enrichment with COSMOS.cA549 cells spiked into healthy donor whole blood at concentrations of 400 cells/mL or 40 cells/mL and processed and sorted with COSMOS. An additional CD45 depletion step was used to partly enrich the A549 cells prior to COSMOS sorting. A549 cells were pre-stained with a fluorescent cell tracker dye before spike-in. A portion of cells after RBC lysis and after CD45 depletion were analyzed with flow cytometry to estimate the fraction of A549 cells in the mixtures after each step. A549 cell purities were estimated from SNP analysis of sorted cells.

**Table S4.** Cell viability after flowing through or sorting on COSMOS.

| Sample and cell type | Number of runs | % Viability of pre-sorted cells (average ± SD) | % Viability of sorted or flown-through cells (average ± SD) |
| --- | --- | --- | --- |
| PBMC | 13 | 94.7 ± 2.5% | 94.2 ± 3.8% |
| B-lymphoblastoid cell line (GM12878) | 10 | 90.6 ± 3.1% | 91.0 ± 5.4% |
| Cancer cell lines (H522 and A549) | 13 | 96.1 ± 2.0% | 96.9 ± 2.2% |
| Dissociated NSCLC tissue | 3 | 71.0 ± 4.4% | 65.0 ± 1.0% |

**Table S5.**
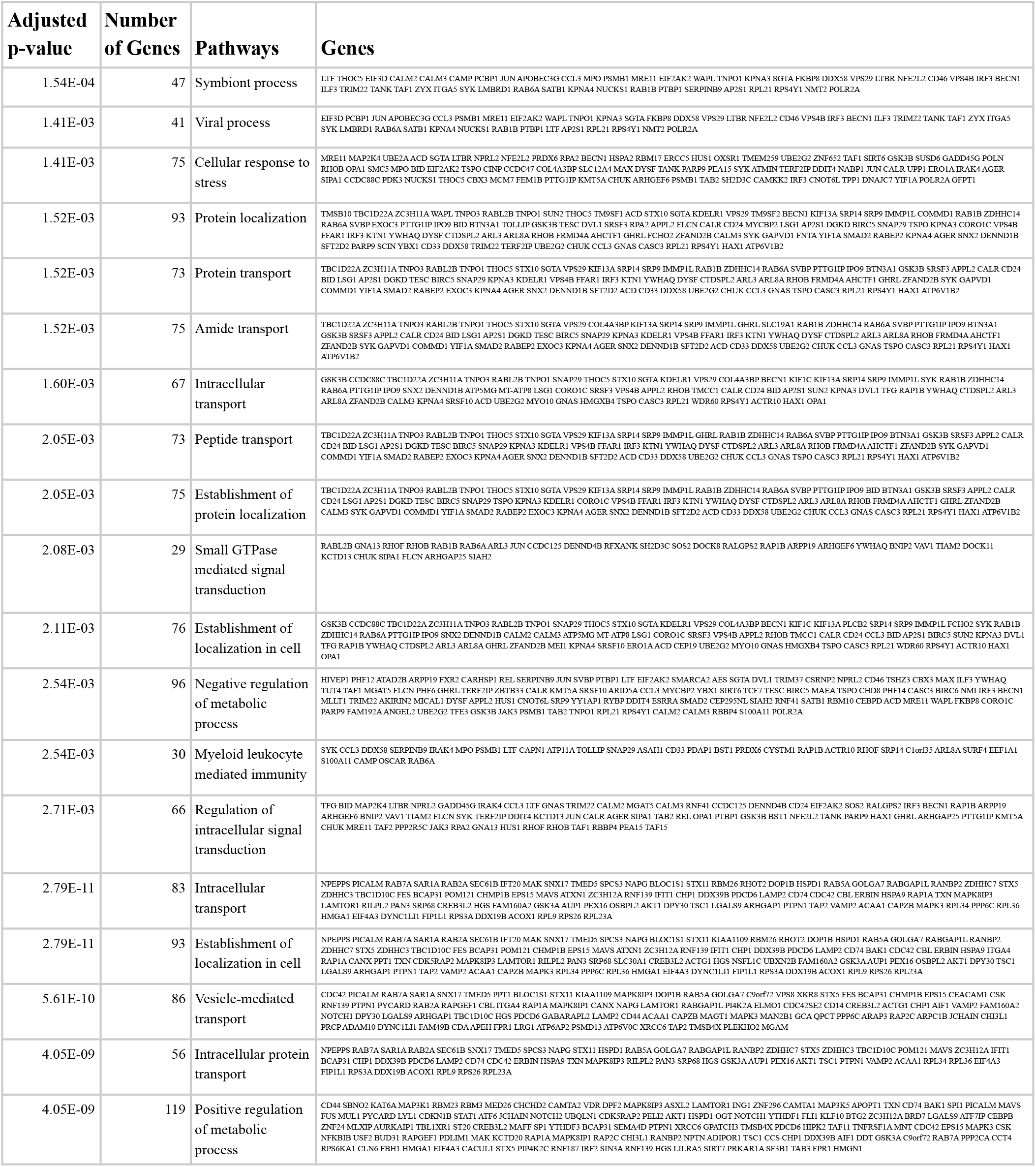
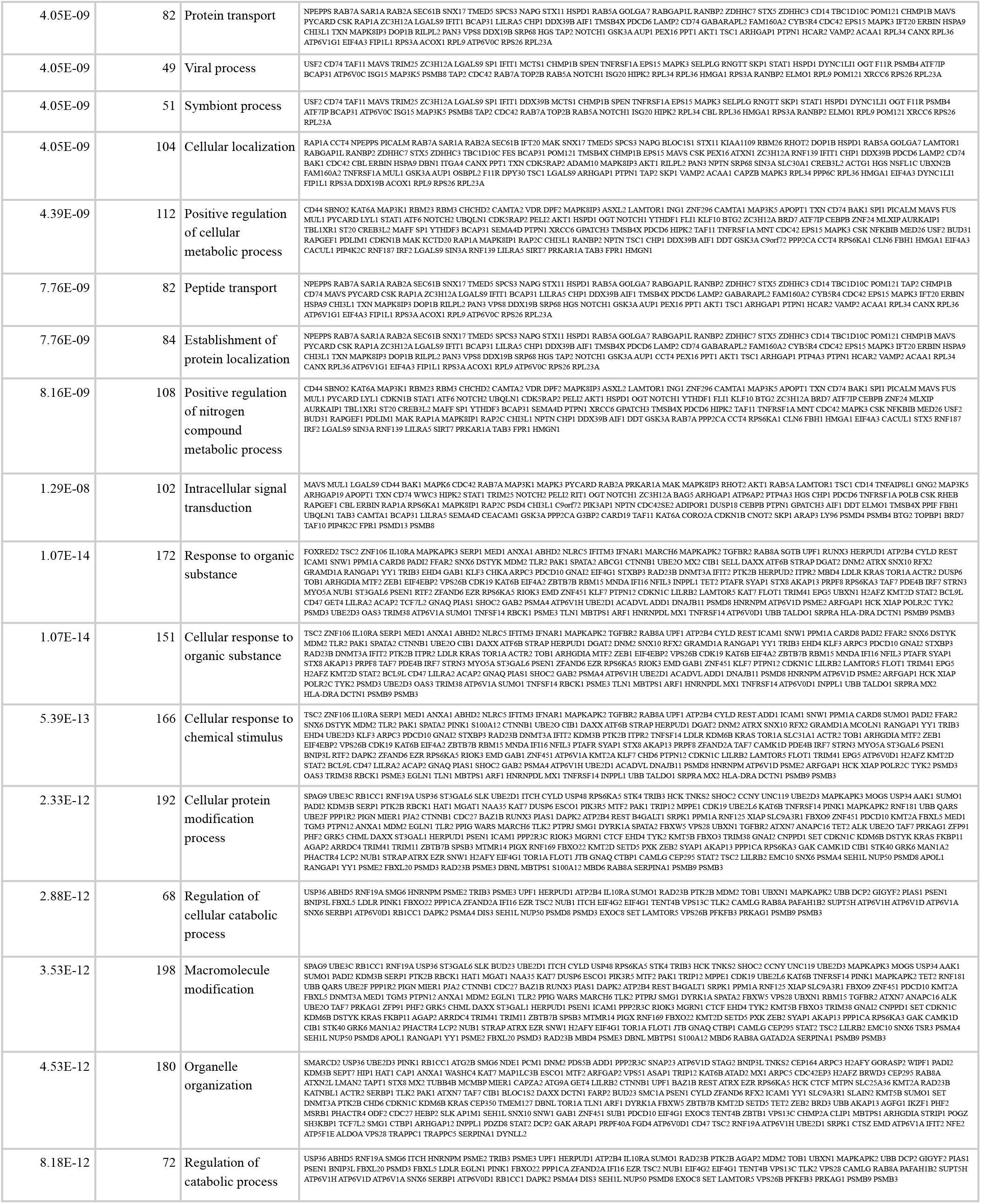
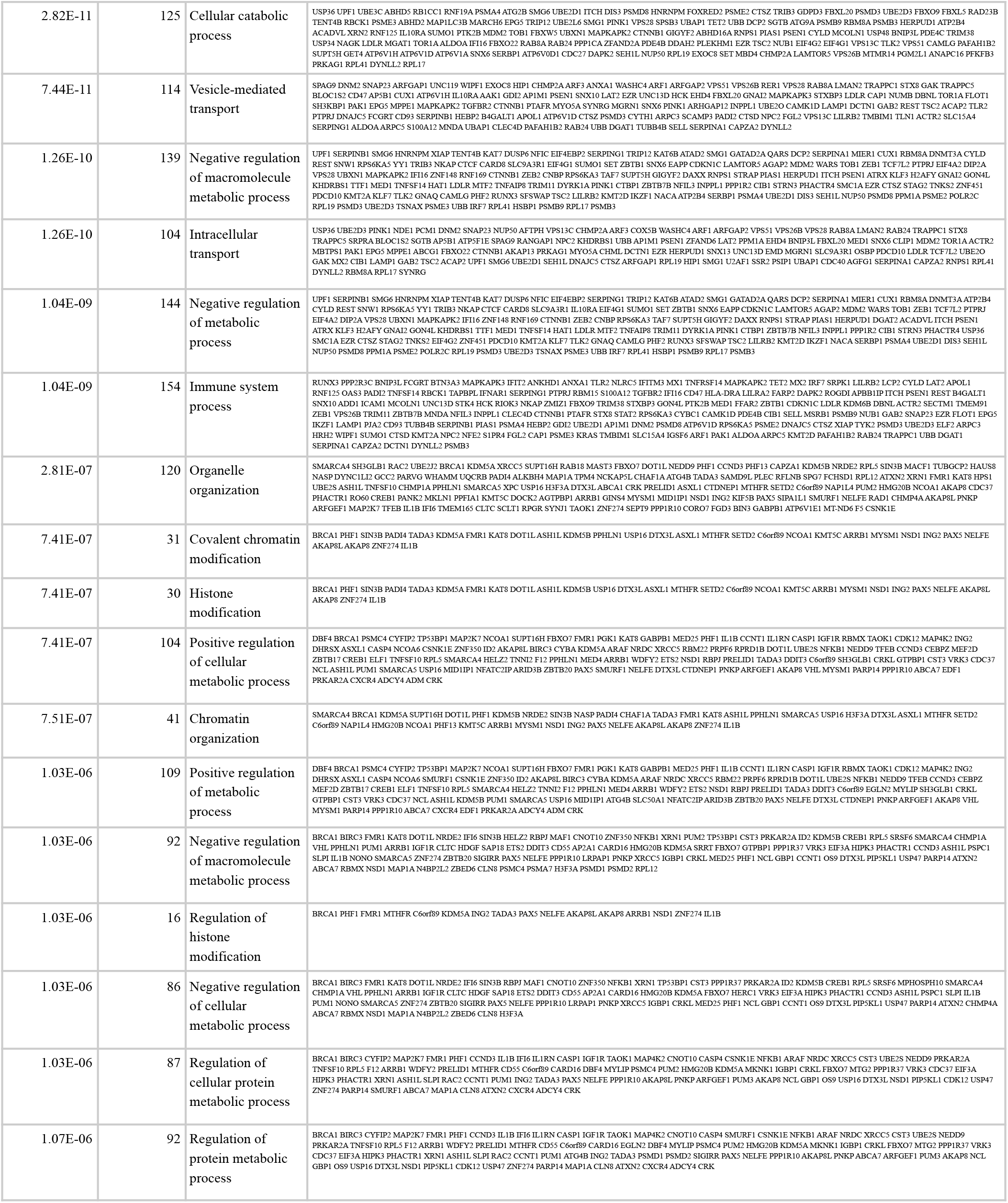
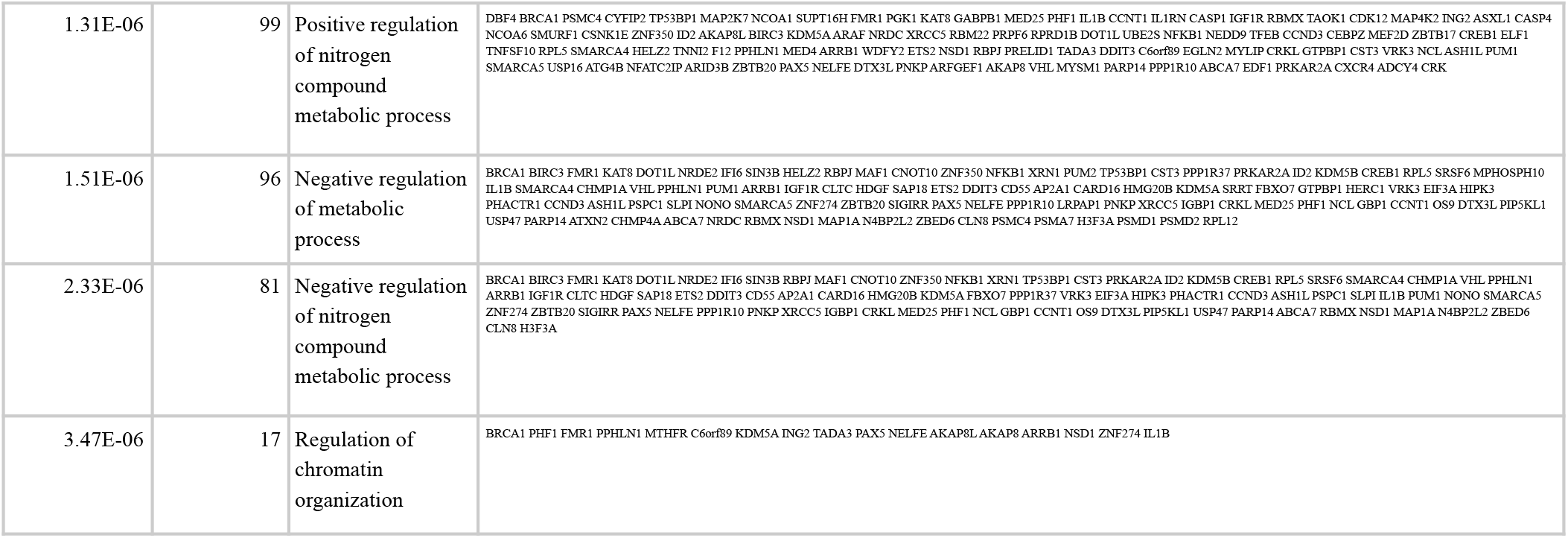
Full list of enriched pathways in FACS-sorted vs COSMOS-sorted neutrophils.

**Figure S1.**
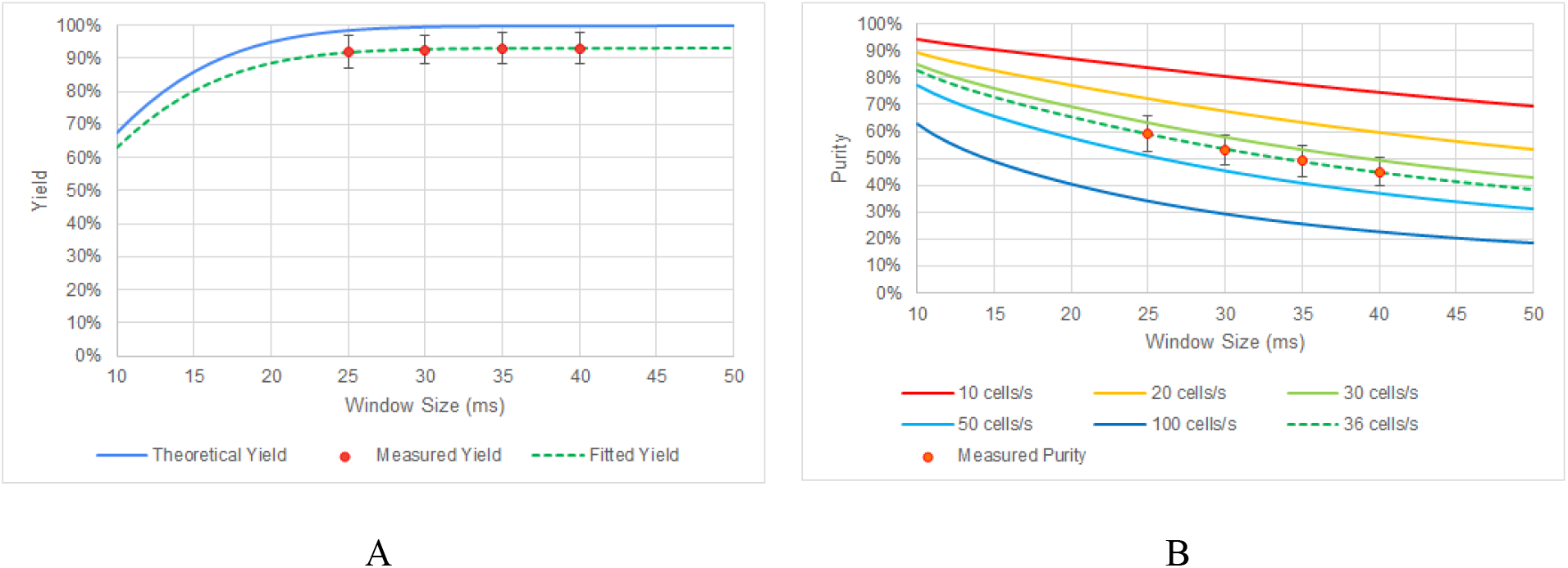
Performance of 0.5% random sorting of PBMC samples using different window sizes (25, 30, 35 and 40 milliseconds). A total of 341 experiments were run across four window sizes in 21 microfluidic devices (three chips each from seven photoresist mold sets) and on two hardware systems. (**A**) Yield: The theoretical curve assumes a normal distribution of cell arrival time with a standard deviation of 5 ms; fitted curve adds a limit of detection level at 93%. (**B**) Purity: Solid and dotted lines are theoretical values at various cell throughput; ±3 ms exclusion zone is assumed around each cell to match measured values with the theoretical values. The error bars in both graphs represent one standard deviation (2σ total) of the raw experimental data in each window size.

**Figure S2.**
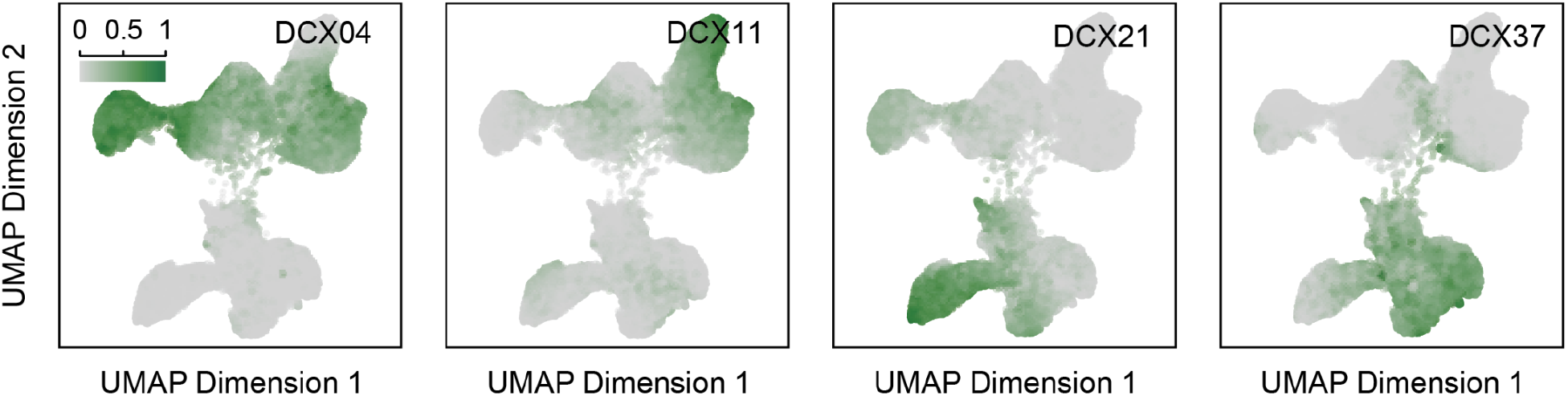
The UMAP projection in (**fig. 2A**), colored by the value of each coordinate in the embedding space of the model, demonstrating the contribution of that coordinate in identifying cells that are highlighted. For example, the leftmost plot demonstrates the value of coordinate number 4 (DCX04) and it shows that this coordinate “encodes” for the NSCLC and HCC (malignant) cells, whereas coordinates DCX11, DCX21 and DCX37 correspond to the HCC, fnRBC and PBMC classes, respectively.

**Figure S3.**
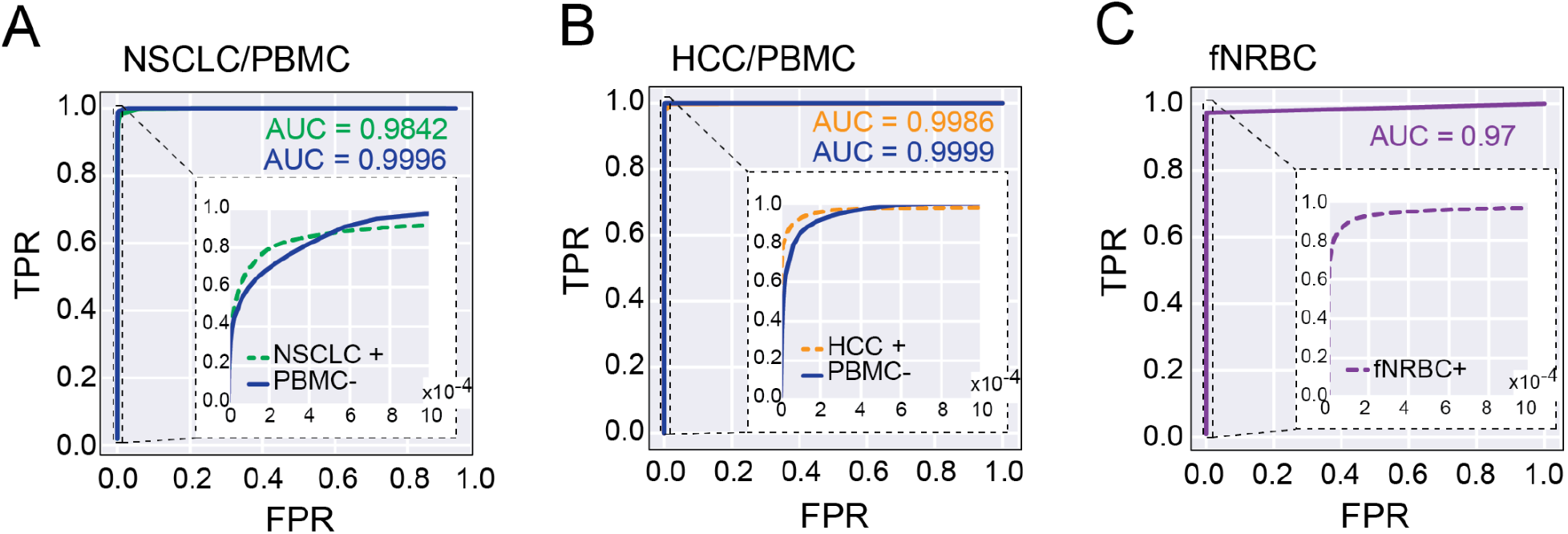
(**A and B**) Receiver operating characteristic (ROC) curves for the classification of (**A)** NSCLCs and (**B**) HCCs. Two ROC curves each are shown: one for the positive selection of each category, and one for negative selection, specifically for the selection of non-blood cells. Area Under Curves (AUCs) achieved for NSCLC are 0.9842 (positive selection) and 0.9996 (negative selection) and for HCC are 0.9986 (positive selection) and 0.9999 (negative selection). (**C**) ROC curves for the classification of fnRBCs and the AUC is 0.97 (positive selection). Insets zoom into the upper left portions of the ROC curves where false positive rates are very low to highlight the differences between modes of classification.

**Figure S4.**
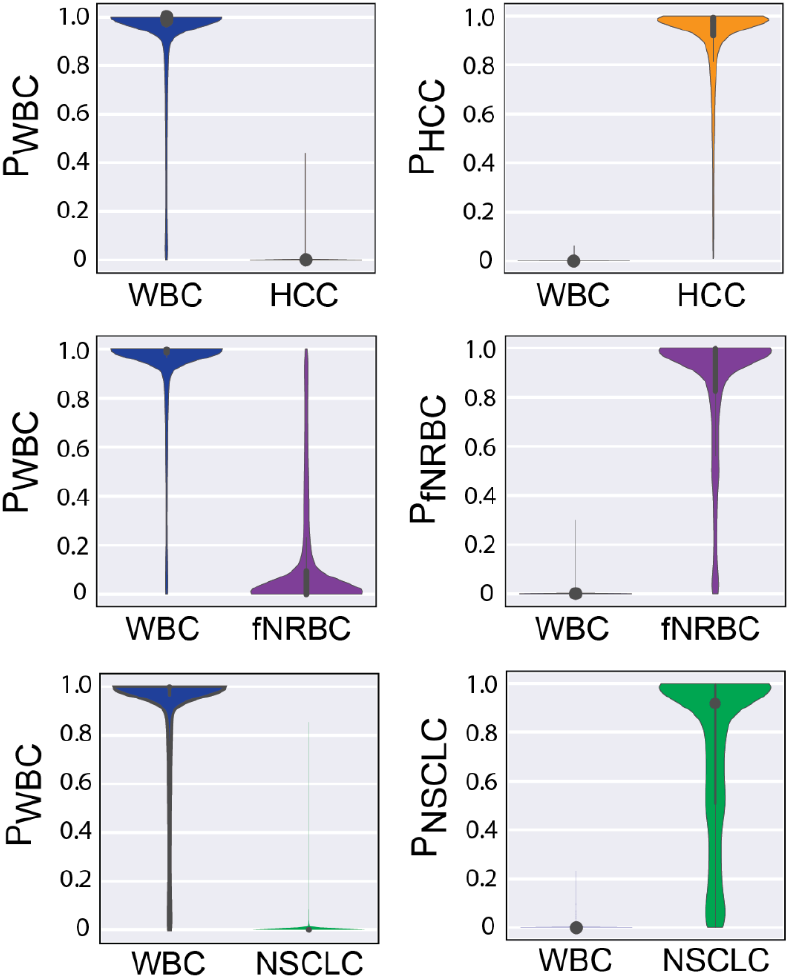
Violin plots showing the predicted probabilities of assigning cells in each category to its appropriate class. The plot on the left shows the probability distribution of PBMCs as well as NSCLCs being classified as PBMCs (P_PBMC_) and the plot on the right shows the probability distribution of PBMCs as well as NSCLCs being classified as NSCLCs (P_NSCLC_).

**Figure S5.**
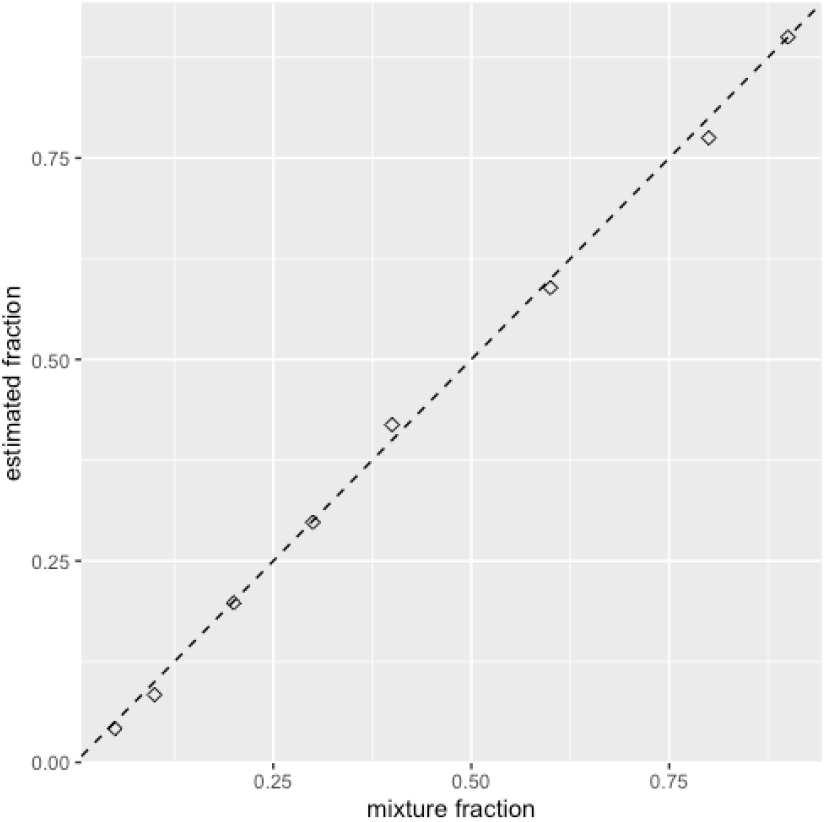
Accuracy of SNP-based mixture fraction estimates in control DNA mixtures. Each composite sample contained 250 pg of bulk DNA drawn from two individuals and the mixture proportion of DNA from the second individual was set at 5%, 10%, 20%, 30%, 40%, 60%, 80% and 90%. A close correspondence was found between the known and estimated mixture proportions.

**Figure S6.**
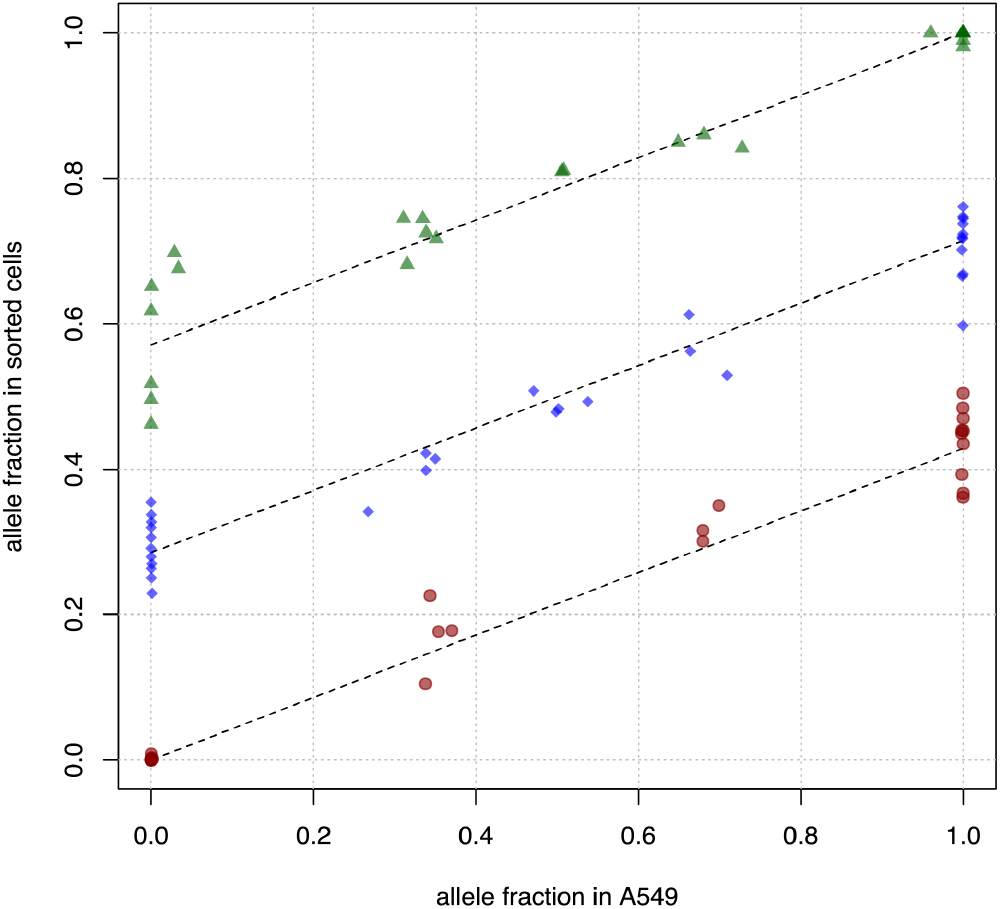
A549 purity in cells enriched using COSMOS from a 40 cells/mL spike-in into healthy donor whole blood. The purity and blood sample genotypes were estimated with an expectation-maximization (EM) algorithm. Green circles, blue diamonds and red triangles denote AA, AB and BB genotypes respectively in the blood sample used as a base for the spike-in mixture; dotted lines represent the expected allele fractions for the three blood genotypes at the inferred purity of 43% (95% confidence interval 0.40 - 0.45) which is also the slope of the lines.

**Figure S7.**
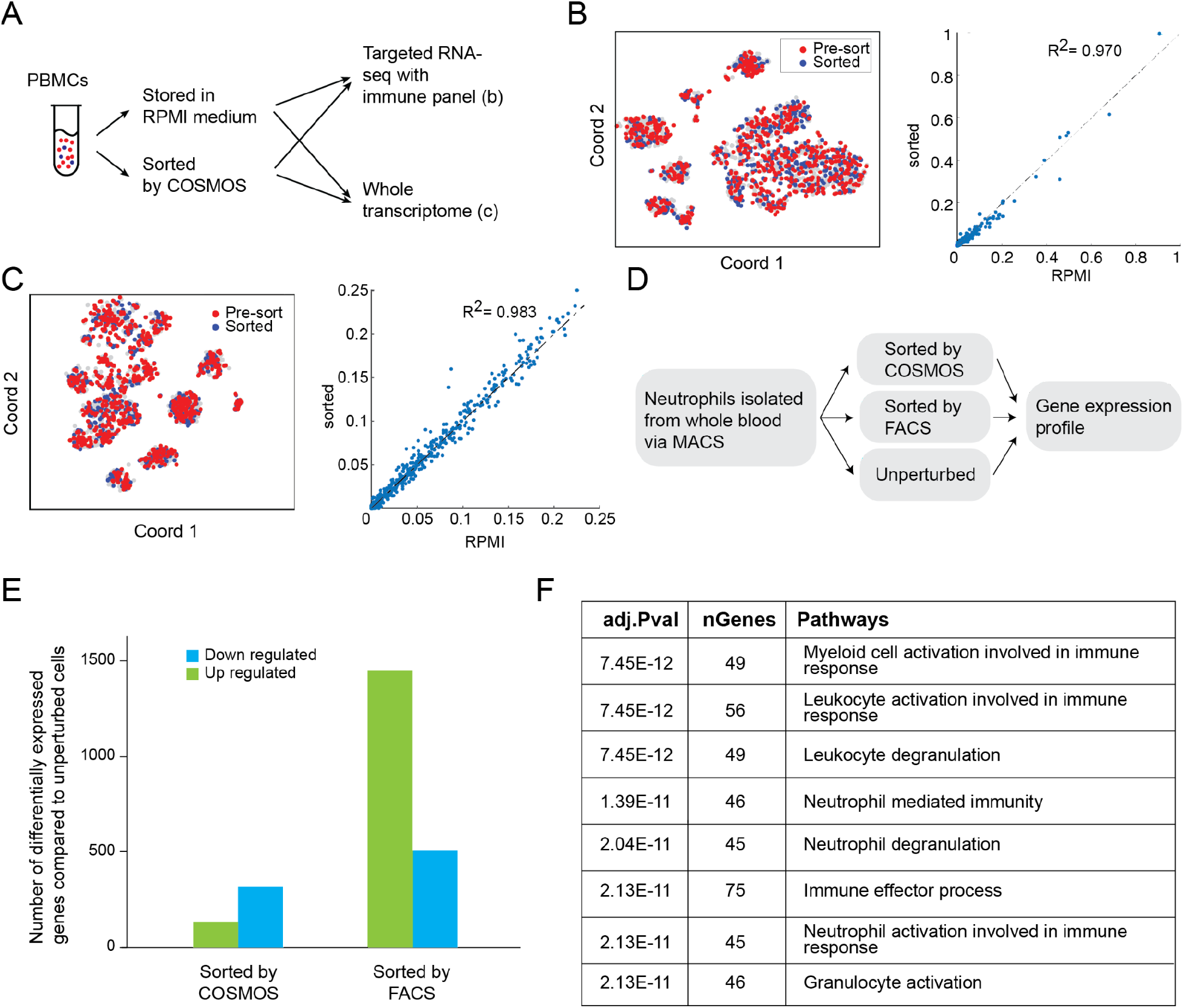
Cell health and quality after COSMOS sorting. (**A**) Workflow schematics: PBMCs flowed-through COSMOS were compared with control cells stored in RPMI medium using either a single cell targeted immune panel (**b**) or single cell whole transcriptome (WTA) workflow (**c**) with the BD Rhapsody™ system. (**B and C**) Left is a t-SNE plot of gene expression profiles of the pre-sort and sorted cells, each point is a cell. Right is a correlation plot of mean (log_10_(molecules per cell per gene)) for the two conditions, each point is a gene. The two samples overlapped with each other in t-SNE plot and gene expression levels showed high correlations (R^2^ equaled 0.97 and 0.98 respectively for targeted panel and WTA), indicating no significant gene expression change after sorting. (**D-G**) COSMOS sorting of unlabeled neutrophils yielded healthier cells compared to stained and FACS sorted cells. (**D**) Workflow schematics: human neutrophils were first isolated from whole blood by immunomagnetic negative selection then split into multiple aliquots for four conditions: unperturbed, stained and flow-sorted by FACS, unstained/unlabeled and sorted by COSMOS. Pre-sorted and sorted cells were lysed for bulk gene expression profiling by RNAseq. (**E and F**) Bulk RNAseq gene expression analysis of the cells in different groups. (**E**) Number of up- and down-regulated genes compared to unperturbed cells, confirmed that COSMOS-sorted cells had minimal gene expression differences compared to unperturbed cells, much fewer than FACS-sorted cells did. (**F**) Upregulated pathways in FACS-sorted cells compared to COSMOS-sorted cells, suggests that FACS induced upregulation of pathways in neutrophil activation and degranulation.

**Figure S8.**
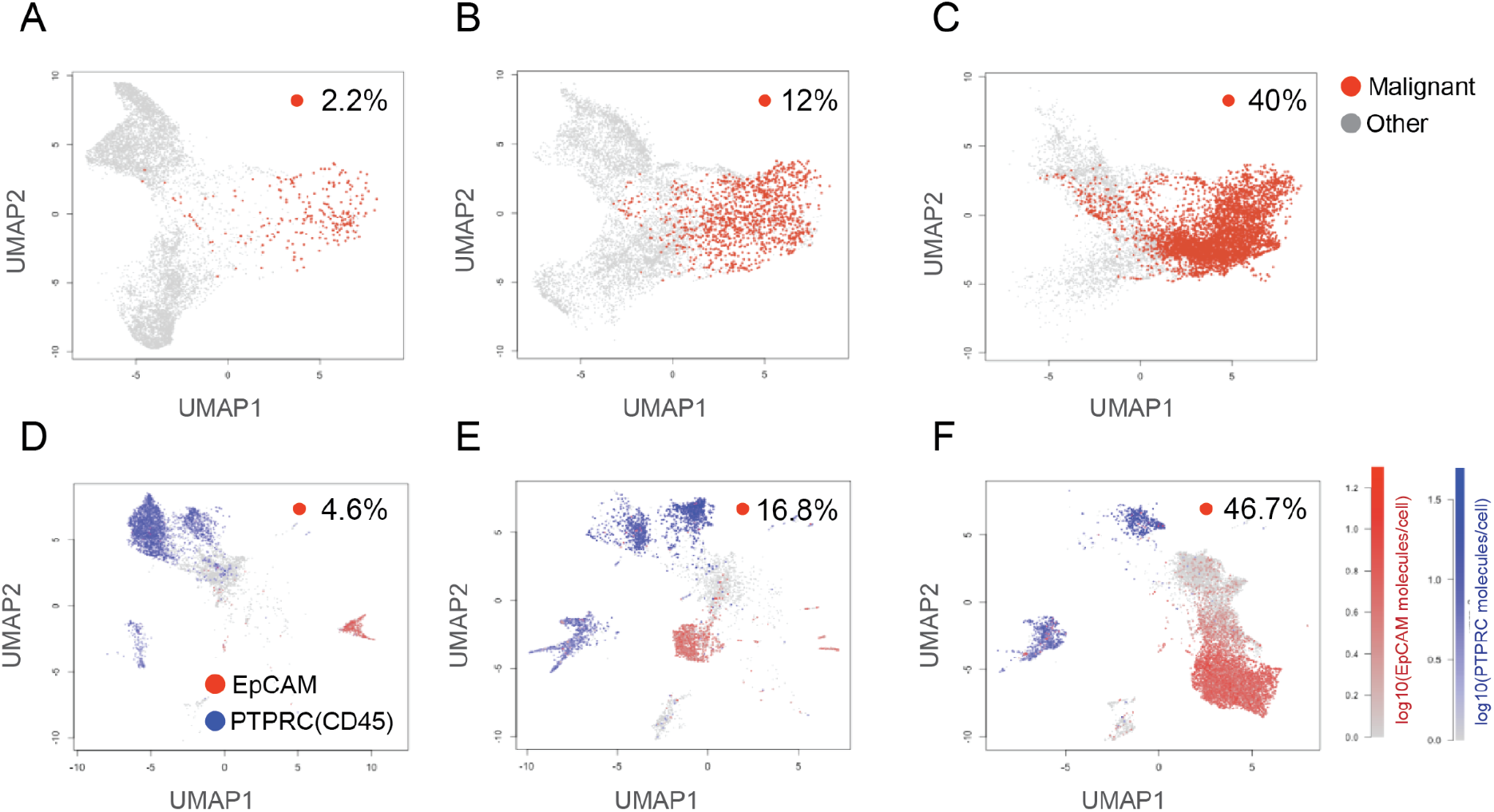
UMAP of morphology embeddings vs scRNAseq gene expression of three different dissociated tumor cell (DTC) samples. Samples from patients with lung adenocarcinoma containing low (**A, D**), medium (**B, E**) and high (**C, F**) percentage of malignant cells were tested. (**A-C**) UMAP of morphological embeddings: each data point is a cell; the predicted malignant cells are colored red and non-malignant cells colored gray. The plot labels indicate the fraction of malignant cells predicted by the model. (**D-E**) UMAP of single cell RNA gene expression profiles from all genes. The red and blue color gradients indicate the expression levels of EpCAM (tumor cell marker) and PTPRC (CD45, immune cell marker) respectively. The plot labels indicate the fraction of EpCAM+/PTRPC-cells. Overall, the morphology-based model predicted a similar fraction of malignant vs nonmalignant cells and UMAPs have similar resolution and separation of malignant vs nonmalignant cells for all three samples.

**Figure S9.**
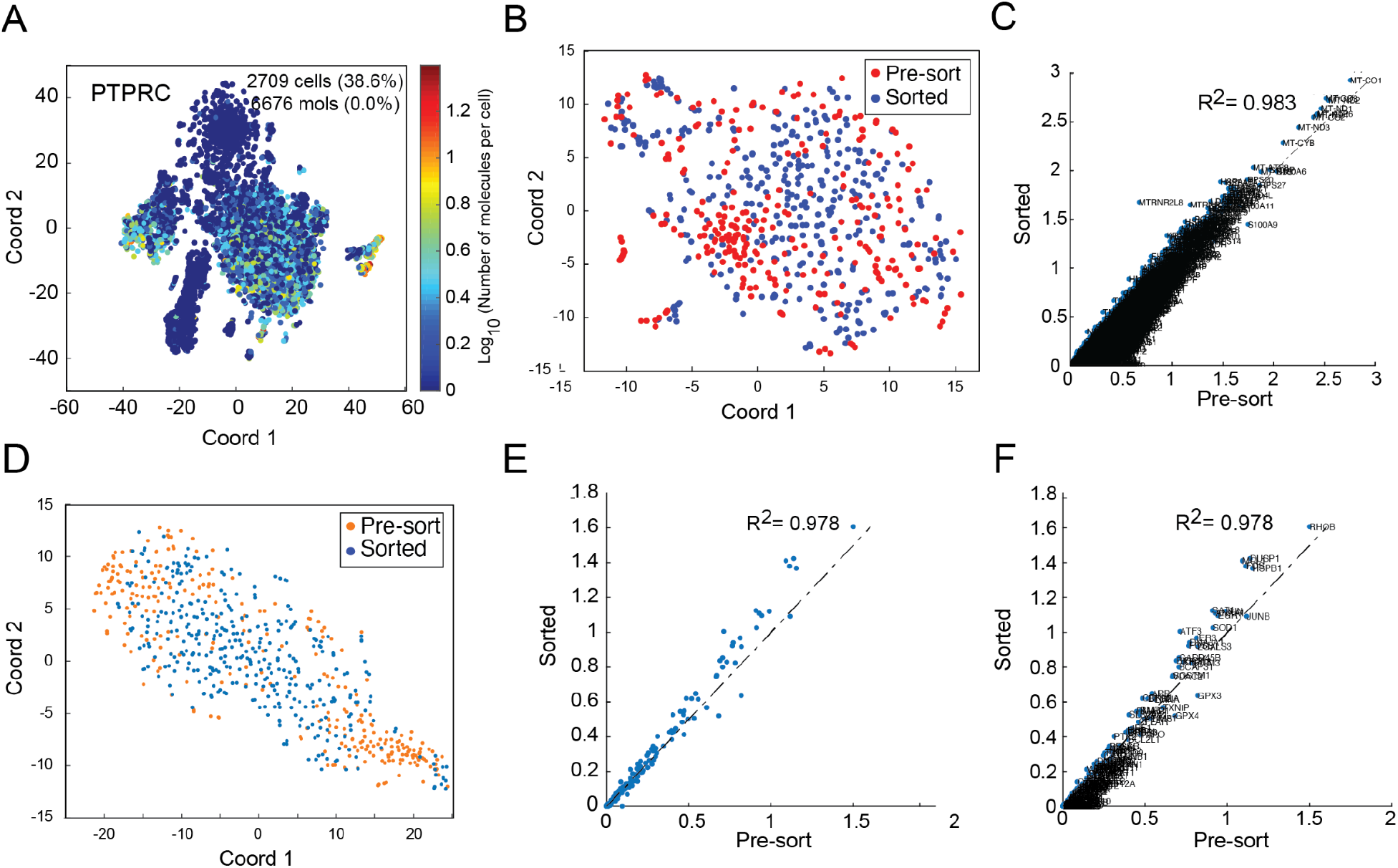
(**A**) Pseudo-color gene expression level of PTPRC (CD45, immune cell marker) in the DTC t-SNE plot. In combination with **fig. 3J** it confirmed that the Sorted cells were mostly in the EPCAM^+^/PTPRC(CD45)^-^ cluster. (**B**) Further sub-clustering of the EPCAM^+^/PTPRC(CD45)^-^ cluster showed that sorted cells almost completely overlapped with pre-sorted cells for all subclusters and (**C**) the gene expression profiles are highly correlated (gene correlation plot, each dot is a gene with the gene names annotated). (**D-F**) Stress and apoptosis related gene expression profile comparison of the pre-sorted and sorted cells in the EPCAM+/CD45-subpopulation. (**D**) t-SNE plot of the EpCAM^+^/CD45^-^ cluster from **fig. 3I** using only the 166 stress and apoptosis genes, showing sorted cells overlap with pre-sort cells in all subclusters. (**E and F**) Gene expression profiles were highly corrected between sorted and pre-sort EpCAM+/CD45 cells; the correlation coefficient was 0.978 (each data point is a gene, **c** has gene names annotated), suggesting that COSMOS sorting did not cause additional cell stress.

## Acknowledgements

We thank Catherine Blish for input and suggestions on the neutrophil study design and the manuscript and Prashast Knandelwal and Tariq Sharaat for early contributions to the COSMOS system development.

## Funding

This work was supported by private funding.

## Author contributions

Research design and study concept: MS, MM, NL, HPC, EAA, KPP, TJM

Design studies: MS, MM, NL, HPC, KPP, KBJ, KS, AJ, EJL, CC, PN, SH

Perform experiments and analysis: KS, AJ, EJL, CC, PN, SH, RC, JM, KPP, AYWT, QFS, JC, JW, BC

Figures: MM, KPP, KBJ, CC, KS

Drafted manuscript: MM, MS, KBJ, NL, TJM, KS, AJ, CC, HPC, KPP, CJ

Review & editing manuscript: all authors

## Competing interests

All authors are current or former employees at or are affiliated with Deepcell Inc.

